# Ratiometric sensing of Pnt and Yan transcription factor levels confers ultrasensitivity to photoreceptor fate transitions in Drosophila

**DOI:** 10.1101/430744

**Authors:** Sebastian M. Bernasek, Suzy SJ Hur, Nicolás Peláez-Restrepo, Jean-François Boisclair Lachance, Rachael Bakker, Heliodoro Tejedor Navarro, Nicelio Sanchez-Luege, Luís A. N. Amaral, Neda Bagheri, Ilaria Rebay, Richard W. Carthew

## Abstract

Cell state transitions are often triggered by large changes in the absolute concentrations of transcription factors and therefore large differences in the stoichiometric ratios between these factors. Whether cells can elicit state transitions using modest changes in the relative ratios of co-expressed factors is unclear. In this study we investigate how cells in the *Drosophila* eye resolve cell state transitions by quantifying the expression dynamics of the ETS transcription factors Pnt and Yan. We find that eye progenitor cells maintain a relatively constant ratio of Pnt/Yan protein despite expressing both proteins with pulsatile dynamics. A rapid and sustained two-fold increase in the Pnt/Yan ratio accompanies transitions to photoreceptor fates. Genetic perturbations that modestly disrupt the Pnt/Yan ratio produce fate transition defects consistent with the hypothesis that transitions are normally driven by a two-fold shift in the ratio. A biophysical model based on cooperative Yan-DNA binding coupled with non-cooperative Pnt-DNA binding illustrates how two-fold ratio changes could generate ultrasensitive changes in target gene transcription to drive fate transitions. In this way, coupling cell state transitions to the Pnt/Yan stoichiometric ratio sensitizes the system to modest fold-changes, conferring both robustness and ultrasensitivity to the developmental program.

## INTRODUCTION

Organismal development proceeds through a sequence of cell transitions that yield increasingly restricted cellular states or fates. A common feature of cells that undergo fate transitions is the apparent irreversibility of the transitions, even when triggered by transient stimuli. Dynamical modeling of these transitions often assumes multistability, in which cells can be resting in one of two or more stable states (Rand et al., 2021; Sáez et al., 2022). These stable states are typically a progenitor state and one or more derived states with lesser developmental potential.

Transcription factors are frequently utilized to define cell states, and transitions to a new state are often driven by varying their absolute or relative protein abundance (Laslo et al., 2006; Park et al., 2012; Yao et al., 2008). A considerable number of developmental transitions rely on mutually exclusive expression of two antagonistic transcription factors. Each factor programs a distinct cell state such that cells adopt one of two possible fates depending on whether they express high levels of one transcription factor or the other. Transitions are triggered when transient stimuli alter the balance between the two factors, resulting in expression of a new dominant factor. First documented in the early embryo of *Drosophila* (Jaeger et al., 2004), such regulatory switches built on mutual repression function in mouse embryonic germ-layer restriction, hematopoiesis, neural tube patterning, mesoderm development, and pancreatic lineage restriction (Acloque et al., 2011; Briscoe and Ericson, 2001; Kumar et al., 2017; Lagha et al., 2009; Schaffer et al., 2010; Schrode et al., 2014). A hallmark of such systems is the mutually exclusive expression of the two opposing transcription factor regulators, which is frequently controlled by a bistable mechanism and double negative feedback loops connecting the two factors.

In *Drosophila*, two ETS-domain transcription factors, Pointed (Pnt) and Yan (also known as Aop, Anterior open), act downstream of signals mediated by receptor tyrosine kinases (RTKs) to regulate cell fate transitions in a wide variety of tissues across the body and across the life cycle (Flores et al., 2000; Gabay et al., 1996a; Gabay et al., 1997; Halfon et al., 2000; Lachance et al., 2014; Melen et al., 2005a; O’Neill et al., 1994; Rebay and Rubin, 1995; Rogge et al., 1995; Shwartz et al., 2013; Xu et al., 2000). Consistently across many different developmental transitions, loss of *yan* results in too many progenitors transitioning to a particular fate, while loss of *pnt* prevents fate transitions (Brunner et al., 1994; Flores et al., 2000; Gabay et al., 1996a; Halfon et al., 2000; O’Neill et al., 1994; Xu et al., 2000). Since both Pnt and Yan bind to a common GGA(A/T)-containing DNA sequence motif (Nitta et al., 2015; Wei et al., 2010; Zhu et al., 2011), competition for DNA occupancy and antagonistic regulation of common target genes is thought to orchestrate the transcriptional changes that drive particular transitions (Flores et al., 2000; Halfon et al., 2000; Hollenhorst et al., 2011; Lachance et al., 2018; Nitta et al., 2015; Webber et al., 2013a; Webber et al., 2013b; Webber et al., 2018; Wei et al., 2010; Xu et al., 2000; Zhu et al., 2011).

The opposing genetic phenotypes and transcriptional activities of Pnt and Yan, together with their mutually exclusive expression patterns in certain tissues, inspired development of a bistable model for cell fate specification based on mutual repression of one another’s expression (Gabay et al., 1996b; Golembo et al., 1996a; Graham et al., 2010; Melen et al., 2005b). For example, in the embryonic ventral ectoderm, ventral-most cells express Pnt while ventro-lateral cells express Yan (Gabay et al., 1996b). This pattern is established by secretion of a ligand for the EGF Receptor (EGFR) from the ventral midline (Golembo et al., 1996b). EGFR activation in nearby cells induces the Ras pathway to express Pnt and degrade Yan (Gabay et al., 1996b; Melen et al., 2005b). Cells with insufficient EGFR activation express Yan, which represses Pnt. Dynamical modeling has described this as a bistable system in which cells assume different fates based on whether they are in a low Pnt/high Yan state vs. a high Pnt/low Yan state (Graham et al., 2010; Melen et al., 2005b).

While the bistable model readily explains transitions in which the different states are marked by mutually exclusive Pnt and Yan expression, an unresolved paradox is that many tissues have extensive co-expression of Yan and Pnt (Lachance et al., 2014). The third-instar larval eye disc is one such tissue. Qualitative imaging has shown that rather than adopting mutually exclusive expression states, eye progenitor cells co-express high levels of Pnt and Yan for prolonged periods. Further, when progenitors transit to photoreceptor (R cell) fates, they continue to co-express both proteins, transiently increasing Pnt levels before initiating a parallel decay in both Pnt and Yan (Lachance et al., 2014). These observations are at odds with the long-standing assumption that cross-repression within a bistable network creates mutually exclusive Yan and Pnt expression states during R cell fate specification (Graham et al., 2010; Lai and Rubin, 1992; Rebay and Rubin, 1995).

Here, we have explored how Yan and Pnt expression dynamics in the third instar eye disc impact progenitor to R-cell fate transitions. Quantitative fluorescence-based microscopy was coupled with automated high-throughput image analysis to measure Yan and Pnt protein levels at single cell resolution in thousands of eye disc cells. We find that despite variation in Pnt and Yan levels, cells in the progenitor state maintain a fairly constant ratio of Pnt/Yan for up to 50 hours of development. When specific progenitors transit to R-cell fates, the Pnt/Yan ratio rapidly increases, but only by at most two-fold. Experimental perturbation of the ratio results in progenitor cells either prematurely transiting or failing to transit to R-cell fates, implicating the Pnt/Yan ratio as the driver of cell fate transitions. Using a biophysical model, we demonstrate how even a modest change in the Pnt/Yan ratio will cause a large change in the DNA occupancy of the two proteins on their common target genes. We conclude that the transition to photoreceptor fates is sensitive to the stoichiometric ratio of Pnt and Yan and propose that ratiometric sensing mechanisms could confer ultrasensitivity to biological transitions regulated by multiple factors.

## RESULTS

During the final 50 hours of the larval phase of the *Drosophila* life cycle, a morphogenetic furrow (MF) moves across the eye disc epithelium at a constant velocity from posterior to anterior (Figure 1A). As the MF moves, every 2 hours a column of periodic R8 photoreceptor cells is formed with near simultaneity immediately posterior to the MF. Each R8 cell in the column secretes ligands that activate the RTK – Ras – MAPK pathway in seven neighboring progenitor cells and causes them to transit from progenitor to R-cell fates (Freeman, 1996). These transitions occur stepwise every ∼ 2 hours, first generating R2 and R5 cells, then R3 and R4 cells, then R1 and R6 cells, and finally the R7 cell (Figure S1A; (Wolff and Ready, 1993)). Since the MF repeatedly induces new columns of R-cell clusters or ommatidia, it ultimately forms a lattice pattern of ∼750 ommatidia in each eye. Although ∼6,000 progenitor cells adopt R-cell fates, a pool of ∼14,000 progenitors remain uncommitted, and later adopt other retinal cell fates (Wolff and Ready, 1991).

**Figure 1.**
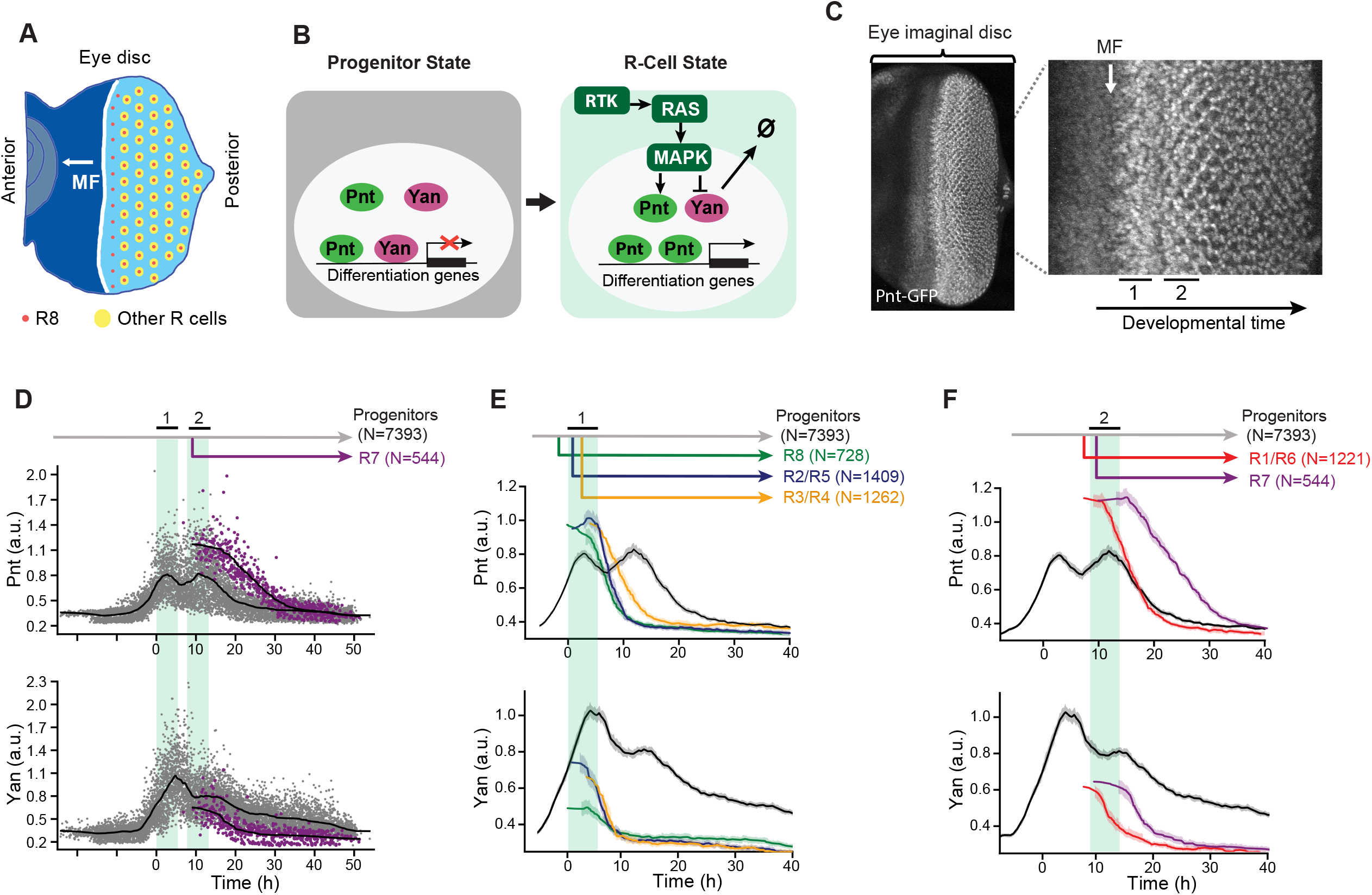
Pnt and Yan expression defines two waves of cell state transitions. **(A)** Schematic of a developing eye disc dissected from a larva ∼100 hours post fertilization. R cell fate specification is initiated by passage of the MF, which moves across the eye disc from posterior to anterior (white arrow). Progenitor cells anterior to the MF (dark blue) do not undergo lineage restriction, while progenitor cells posterior to the MF (light blue) become competent to differentiate into R cells. Specification of regularly spaced R8 cells (red dots) marks the start of ommatidial assembly and is followed by recruitment of additional R-cell types (yellow dots). Adapted from (Peláez et al., 2015). **(B)** Schematic summary of RTK-mediated regulation of progenitor to R-cell fate transitions. Progenitor cells co-express Pnt and Yan (left panel). In response to RTK signals, cells increase Pnt and decrease Yan expression (right panel). **(C)** Maximum intensity projection of Pnt-GFP fluorescence in an eye disc oriented anterior to the left and dorsal up. Right panel shows a magnified view of the region marked by dashed white lines in the left panel. The MF (white arrow) is anterior to the zones where progenitor cells undergo fate transitions (black bars, labeled 1 and 2) that coincide with two successive peaks of Pnt-GFP expression. **(D)** Pnt-GFP and Yan protein levels in individual progenitor (grey) and R7 cells (purple) over developmental time. The numbers marked by N refer to the number of cells analyzed in each group. Progenitor cells are present across all time (grey arrow) while R7 cells arise later in time (purple arrow). Solid black lines are smoothed moving averages across 250 and 75 individual nuclei for progenitor and R7 cells, respectively. Black bars labeled 1 and 2 denote the two peaks of Pnt-GFP in progenitor cells. Shaded vertical stripes highlight these two regions, which coincide with transition of progenitor to various R fates. **(E)** Line averages for Pnt-GFP and Yan protein levels in progenitor (grey), R8 (green), R2/R5 (blue) and R3/R4 (orange) cells over developmental time. The numbers marked by N refer to the number of cells analyzed in each group. For all line averages, the 95% confidence intervals are shaded. **(F)** Line averages for Pnt-GFP and Yan protein levels in progenitor (grey), R1/R6 (red), and R7 (purple) cells over developmental time. The numbers marked by N refer to the number of cells analyzed in each group. For all line averages, the 95% confidence intervals are shaded.

Progenitor cells posterior to the MF co-express Pnt and Yan and are poised to adopt R-cell fates (Figure 1B). When progenitors transition to R-cell fates, RTK – Ras – MAPK signals regulate the opposing transcriptional inputs from Pnt and Yan on their shared target genes. The signals concomitantly downregulate Yan activity and upregulate Pnt activity, which is thought to shift the state of the progenitor cell towards an R-cell state by reprogramming target gene expression.

To measure Pnt protein abundance in individual eye cells, we used a genomic transgene of the *pnt* locus in which fast-fold green fluorescent protein (GFP) was inserted in-frame at the carboxy-terminus of the coding sequence, tagging all Pnt isoforms. The *Pnt-GFP* transgene fully complemented null mutations in the endogenous *pnt* gene and completely restored normal eye development (Lachance et al., 2014), demonstrating that the GFP tag does not compromise Pnt function and that all essential genomic regulatory sequences are included. To measure Yan protein abundance, we used a monoclonal antibody that elicits immunofluorescence linearly proportional to native Yan protein concentration (Peláez et al., 2015). This enabled us to measure nuclear Pnt-GFP and Yan protein abundance simultaneously in individual cells. Using an established computational pipeline (Peláez et al., 2015) we segmented all eye cell nuclei and then calculated Pnt-GFP and Yan fluorescence levels and exact 3D positions in each eye disc (Figure S1B-E). A unique feature of eye development allowed us to map each cell’s spatial position in the plane of the eye disc onto developmental time. Because the MF moves at a fixed velocity, forming one column of R8 cells every 2 hours (Basler and Hafen, 1989; Campos-Ortega and Hofbauer, 1977; Gallagher et al., 2022), the distance between the MF and any cell posterior to it is linearly proportional to the time elapsed since the MF passed. This mapping was applied to the thousands of segmented nuclei in each eye disc and repeated for multiple eye discs.

A second feature of eye development enabled us to annotate cell states without relying on marker genes but instead on reproducible changes in cell morphology and apical-basal nuclear position. These stereotyped morphological changes are observable before the earliest known marker genes begin to be expressed, allowing accurate annotation of progenitor and R-cell states (Figure S1A,F) (Peláez et al., 2015; Tomlinson and Ready, 1987; Wolff and Ready, 1993). Cross-referencing with cell-type-specific markers confirmed that we could make cell state assigments with >97% accuracy using only morphology (Table S1). Combining these state assignments with the measurements of Pnt-GFP expression, Yan expression, and developmental time for many thousands of cells generated a composite picture of Yan and Pnt expression dynamics (Figure 1D). Although this approach cannot track the behavior of any individual cell over time, it nevertheless provides a dynamic view of eye cells as they develop. From this composite picture, individual cell behaviors can be inferred.

Focusing first on progenitor cells, we found that both Pnt-GFP and Yan expression dramatically increased coincident with passage of the MF (Figure 1C,D). This rapid increase culminated in two successive peaks of Pnt-GFP expression and a single peak of Yan expression at a time period in between the two Pnt-GFP peaks (Figure S2). The peaks of Pnt-GFP in progenitor cells coincided with the time periods when they transition to R-cell fates (Figure 1C-F). Transitions to R8, R2/R5, and R3/R4 fates occurred during the first peak of expression (Figure 1E), while transitions to R1/R6, and R7 fates occurred during the second peak (Figure 1F). After their peaks, both Pnt-GFP and Yan decayed to a basal level in progenitors (Figure 1D). Despite the alternating peaks of Pnt and Yan expression, the overall trend of induction and decay was highly similar for the two proteins.

Focusing next on Pnt-GFP and Yan dynamics in cells that had transitioned to R-cell fates, we found that these transitions coincided with a rapid increase in Pnt-GFP and rapid decrease in Yan (Figure 1D-F). The Pnt-GFP increase was so rapid that the youngest cells we could classify as R-cells already had a 25-50% higher level of Pnt-GFP than progenitor cells at comparable time points (Figure 1D). Thereafter, the R-cells maintained this elevated Pnt-GFP for 2 – 4 hours before down-regulating Pnt-GFP to a basal level of expression (Figure 1E,F). Similarly, the decrease in Yan in transitioning cells was so rapid that the youngest cells classified as R-cells already had a 25-50% lower level of Yan than progenitors at comparable time points (Figure 1D). Thereafter, R-cells also maintained this diminished level of Yan for 2 – 4 hours before further down-regulating its expression (Figure 1E,F).

Two key conclusions emerged from this analysis. First, Pnt and Yan expression appear positively correlated in progenitor cells, confirming prior qualitative observations (Lachance et al., 2014). Second, transitions to R-cell fates coincide with concomitant but modest up- and down-regulation of Pnt and Yan expression, respectively. This suggests that cell fate specification in the eye does not rely on a classic bistable mechanism in which mutual repression between Pnt and Yan produces large changes in expression to distinguish the two states (Graham et al., 2010; Melen et al., 2005b).

### Pnt and Yan positively regulate each other’s expression

The positive correlation in expression of Yan and Pnt in progenitor cells led us to hypothesize that they might positively regulate each another’s expression. To test this possibility, we first manipulated Pnt protein levels by varying the copy number of the *pnt* gene. If Pnt positively regulates Yan expression, then we predicted Yan levels should change in parallel to altered Pnt levels. Cells with two copies of the *pnt* gene expressed ∼ 50% more Pnt protein than cells with one copy (Figure 2A,B). This difference was observed both in progenitor cells and R cells. In spite of the different levels of Pnt protein in cells with one or two copies of the *pnt* gene, progenitors transited to R-cell fates at similar times (Figure 2A,B), and produced adult eyes with wildtype morphologies (Figure S3). When we measured Yan, we found that Yan levels also differed between the two genotypes, despite both having two copies of the *yan* gene (Figure 2A,B). Yan output was 25-50% greater in progenitor cells with two copies of the *pnt* gene than in cells with one copy, consistent with a role for Pnt in positively regulating Yan expression.

**Figure 2.**
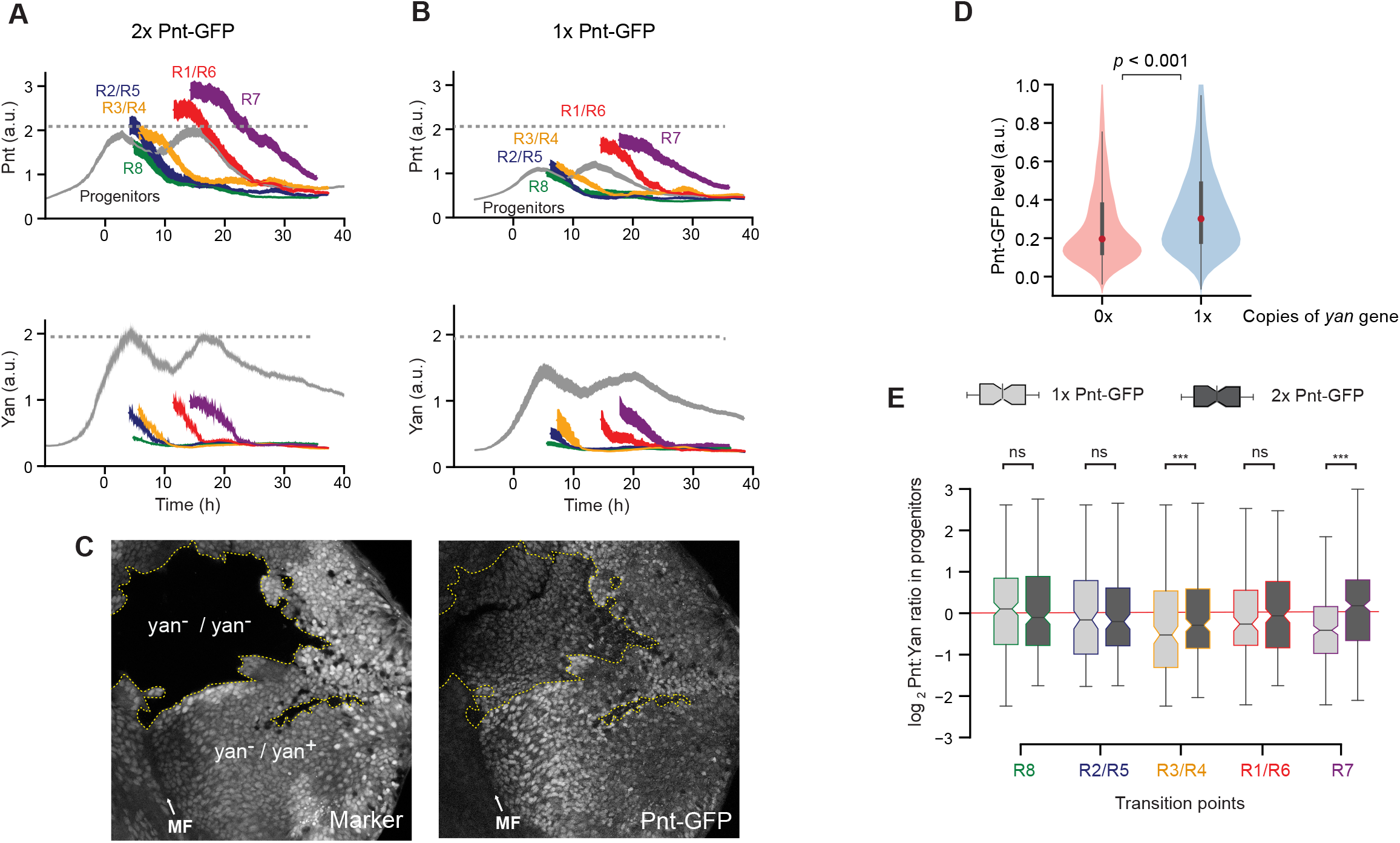
Pnt and Yan activate each other’s expression. **(A)** Line averages for Pnt-GFP and Yan protein levels in progenitor and R cells containing two copies of the *pnt-GFP* gene. Colors marking cell types are consistent with those in Figure 1. Shaded regions are 95% confidence intervals. The number of cells analyzed for each cell-class ranged from 312 (R7 cells) to 3,127 (progenitors). **(B)** Line averages for Pnt-GFP and Yan protein levels in progenitor and R cells containing one copy of the *pnt-GFP* gene. Colors marking cell types are consistent with those in Figure 1. Shaded regions are 95% confidence intervals. The number of cells analyzed for each cell-class ranged from 271 (R7 cells) to 2,834 (progenitors). The horizontal dotted lines in A and B mark the maximal levels of Pnt-GFP and Yan seen in the progenitor cells containing two copies of the *pnt-GFP* gene. **(C)** Confocal section of progenitor cell nuclei in an eye disc containing several patches (clones) of homozygous *yan* mutant cells that are outlined by dotted yellow lines. Left panel, fluorescence from a RFP marker gene that is tightly linked to the *yan* locus. Absence of fluorescence occurs in cells that are missing the RFP marker gene, and therefore are missing the wildtype *yan* gene. Fluorescent cells have one or two copies of the wildtype *yan* gene. Right panel, fluorescence from Pnt-GFP. Note how the fluorescence signal is lower in the *yan* mutant clones. **(D)** Pnt-GFP abundance in homozygous *yan* mutant progenitor cells (0x) versus heterozygous wildtype *yan* progenitor cells (1x). Cells were assigned a *yan* genotype based on measured fluorescence levels of the RFP marker gene linked to *yan*. Violin plots contain red dots that denote the median of each distribution and grey lines that denote the interquartile range. A Mann-Whitney *U* test indicates that the two distributions are significantly different. **(E)** Comparison of the Pnt/Yan ratios in progenitor cells with one copy (light grey filled boxes) versus two copies (dark grey filled boxes) of the *pnt-GFP* gene. Cells were differentially sampled across five time-windows of cell fate transitions, as indicated. These time windows were defined in each disc by the time spanned by the first ten identifiable R cells of a given class. Progenitor cells were sampled from these time windows. Shown are the median ratios boxed by the quartile ratios. The ratio in progenitor cells is statistically indistinguishable between cells bearing different gene dosages during the R8, R2/R5, and R1/R6 cell fate transitions (P>0.1, KS 2-sample test).

To ask if this positive regulatory relationship might be mutual, we next assessed the effect of Yan levels on Pnt-GFP expression by varying the copy number of the *yan* gene. We conducted this experiment by generating clones of *yan* homozygous mutant cells in an eye where other cells have one or two copies of the *yan* gene. Qualitatively, cells with different *yan* gene copy numbers exhibited Pnt-GFP levels that positively correlated with *yan* gene dosage (Figure 2C). Using a published pipeline to segment and measure cells within eye disc clones (Bernasek et al., 2020), we quantified and compared Pnt-GFP protein levels in progenitor cells with zero vs. one wildtype copy of the *yan* gene (Figure 2D). Pnt-GFP expression was ∼30% greater in progenitor cells with one copy of *yan* than in cells with zero copies of *yan* (Figure 2D). This result suggests that Pnt expression is positively dependent on Yan.

The parallel behaviors of Pnt and Yan in these experiments prompted us to quantitatively assess the extent to which Pnt and Yan levels are coordinated within individual cells. To do this, we calculated the Pnt/Yan ratio in progenitor cells containing one or two copies of the *pnt* gene. The Pnt/Yan ratio was invariant to the *pnt* gene copy number in progenitor cells that were sampled at discrete time points over their developmental trajectory (Figure 2E). It was striking that the ratio measured in the progenitors was fairly constant between the sampled time points. To examine this more closely, we calculated the ratio for all progenitor cells over the entire 50 hours of eye development (Figure 3A). The average Pnt/Yan ratio in progenitor cells remained dynamically stable about a fixed value over time (Figure 3A). In conclusion, these results demonstrate that Yan and Pnt expression are tightly and positively coordinated in progenitor cells.

**Figure 3.**
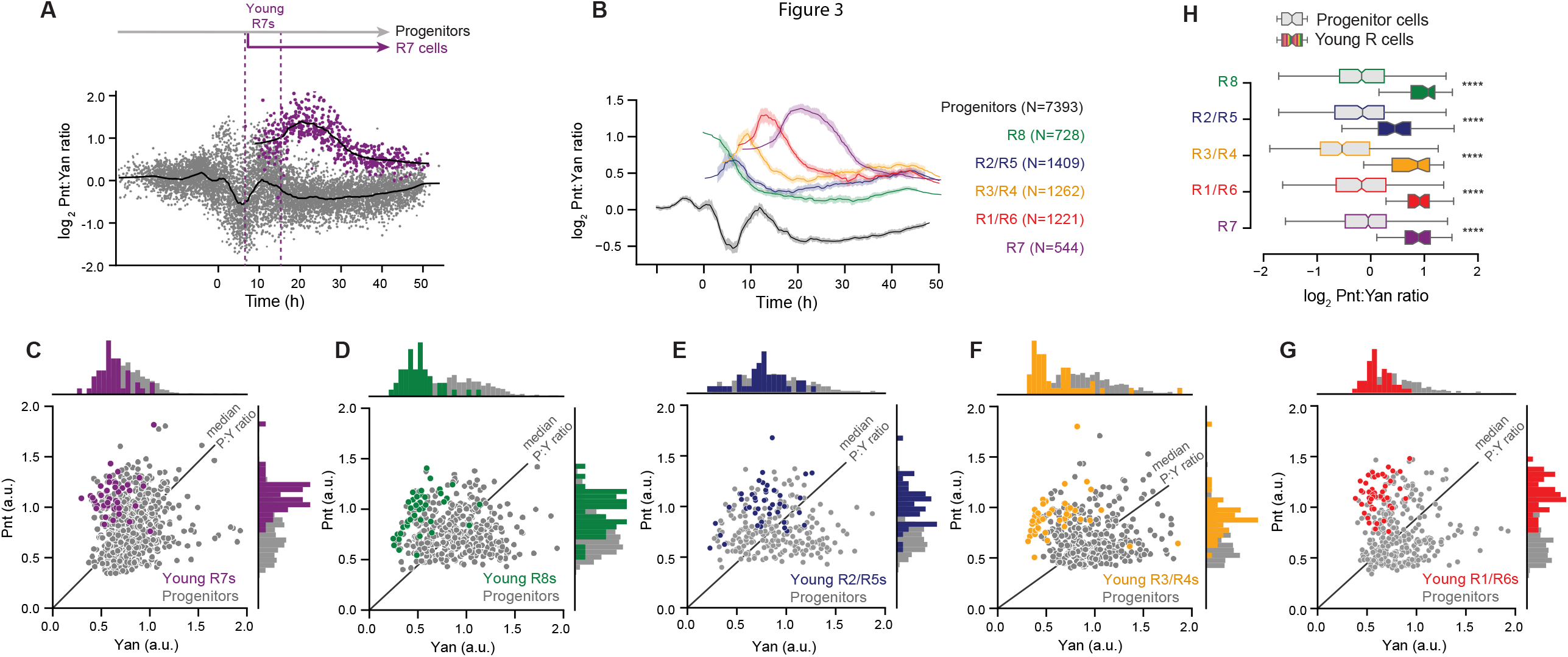
A two-fold shift in the Pnt/Yan ratio accompanies R cell fate transitions. **(A)** The log_2_-transformed Pnt/Yan ratio in individual progenitor (grey) and R7 (purple) cells and their moving line averages (solid black lines). Cells marked as “Young R7” are the initial cohort of R7 cells that are present in the timespan of the first ten identifiable R7 cells. The number of cells analyzed in each class were 7,393 (progenitors) and 544 (R7 cells). **(B)** The line average of the log_2_-transformed Pnt/Yan ratio for all annotated cell types over time. Shading denotes 95% confidence intervals. Colors and N denote cell type and number of cells sampled, respectively. **(C-G)** Joint distributions of Pnt-GFP and Yan protein levels in cells that are Young R7 (C), Young R8 (D), Young R2/R5 (E), Young R3/R4 (F), and Young R1/R6 (G). Colors marking R-cell types are consistent with those colors in Figure 1. R cells were classified as “young” by the criteria described in (A). Progenitor cells (in grey) that were present in the timespan of these young R cells were also analyzed for comparison. Black lines denote the median Pnt/Yan ratios among the progenitor cells. Histograms of the data are shown at the top and side of each scatterplot. Note the overlap between R cells and some of the concurrent progenitors. **(H)** Comparison of the Pnt/Yan ratios in young R cells (color filled boxes and whiskers) versus progenitor cells (grey filled boxes and whiskers) that were sampled over comparable time-periods to the young R cells. A total of 40 cells were analyzed for each group of young R cells, and between 153 and 487 cells were analyzed for each group of progenitors. Shown are the median ratios boxed by the quartile ratios. Asterisks indicate significant differences (P<0.001, KS 2-sample test).

### An increase in the Pnt/Yan ratio accompanies cell state transitions

We next determined the Pnt/Yan ratio for R cells. We first measured the ratio in R7 cells along their developmental trajectory. Transitions to the R7 fate coincided with a rapid increase in the ratio (Figure 3A). The increase was so rapid that the youngest cells we could classify as R7 cells already had a two-fold higher ratio than progenitor cells at comparable time points. Thereafter, the ratio rose and then gradually fell over time, but always remained at higher values than that of progenitor cells. Examination of the Pnt/Yan ratio in all other R-cell types revealed a similar and consistent rapid increase as they transitioned from a progenitor state, followed by a gradual decline to a new steady value that was higher than that of progenitor cells (Figures 3B and S4).

The rapid shift in the Pnt/Yan ratio during cell state transitions suggested that the progenitor ratio becomes destabilized at the onset of these transitions. If so, then cell-to-cell heterogeneity of the ratio should rise during the time-windows of state transitions, and then subside as cell states are resolved. Indeed, greater cell-to-cell variation in the ratio coincided with the times when Pnt and Yan levels peaked (Figure 3A), corresponding to the two waves of R-cell differentiation. To further characterize this instability, we used an established method (Peláez et al., 2015) to quantify the cell-to-cell heterogeneity of Pnt, Yan, and the Pnt/Yan ratio over time (Figure S5). Progenitor cells exhibited maximal ratio heterogeneity during the two time periods that coincide with cell state transitions. Thereafter, heterogeneity diminished in the remaining progenitor cells. R cells exhibited an abrupt drop in heterogeneity immediately after their fate specification, and heterogeneity descended further thereafter. These results support the idea that the Pnt/Yan ratio becomes specifically destabilized when progenitor cells are poised to undergo state transitions.

If cells undergoing state transitions experience a rise in the Pnt/Yan ratio, we should observe cells at transition points that cannot be morphologically classified as R cells but that possess an elevated ratio. This prediction was verified since there was an overlap in ratio values between the youngest cells we could classify as R cells and a subset of other cells sampled at comparable time points (Figure 3C-G). Nevertheless, the median ratio values were significantly different between the R cells and all the other cells (Figure 3H, *P*<0.001, KS 2-sample test). Together these measurements suggest that a subset of progenitor cells elevate their Pnt/Yan ratio by approximately two-fold at transition points, and that this modest increase is sufficient to confer molecular competency for transitioning to a photoreceptor state.

### Perturbation of the Pnt/Yan ratio biases cells towards or against R-cell fates

The previous model for eye development suggested that Yan and Pnt antagonize one another to maintain cells in a progenitor state defined by high Yan and to move cells towards a R-cell state defined by high Pnt (Graham et al., 2010). Our results demonstrate that the cell states are not distinguished by large differences in the Pnt/Yan ratio. Instead, we hypothesize that a modest two-fold change in the ratio is sufficient to trigger R-cell differentiation. We tested this by experimentally altering the ratio in progenitor cells and then asking whether this perturbation altered the normal course of R-cell fate transitions.

Ras-dependent MAPK phosphorylation of Yan and Pnt had been previously found to stimulate turnover of Yan and synthesis of Pnt (Brunner et al., 1994; Gabay et al., 1996a; O’Neill et al., 1994; Rebay and Rubin, 1995). Therefore, we expressed a mutant allele of the *Ras1* gene encoding a constitutively active version of the Ras GTPase, reasoning that it would artificially increase the Pnt/Yan ratio (Figure 4A). Ras^V12^ was transiently expressed in subsets of progenitor cells during the R3/R4 and R7 transition points by driving it with the *sevenless* (*sev*) promoter (Figure S6; (Fortini et al., 1992)). The promoter also drove stable expression in R3/R4 and R7 cells (Figure S6). As predicted, *sev>Ras*^*V12*^ progenitor cells had an elevated second wave of Pnt expression, which coincides with the second time-window of cell state transitions (Figure 4B). Thereafter, progenitors maintained abnormally high levels of Pnt. Yan levels were weakly reduced in progenitor cells during the second transition period (Figure 4C). The overall effect was a modest two-fold elevation in the Pnt/Yan ratio in progenitor cells at the second transition period and thereafter (Figure 4D). When we looked at R-cell fate specification, we observed supernumerary R cells in the *sev>Ras*^*V12*^ eye discs (Figure 4E). These ectopic R cells were typically located beside R7 cells (Figure S7A,B) and had comparable Pnt/Yan ratios to R7 cells (Figure 4E). Thus, just as a two-fold ratio increase accompanies wildtype R7 fate acquisition, artificially inducing a two-fold elevation in the Pnt/Yan ratio causes additional progenitor cells to inappropriately transit to R-cell fates.

**Figure 4.**
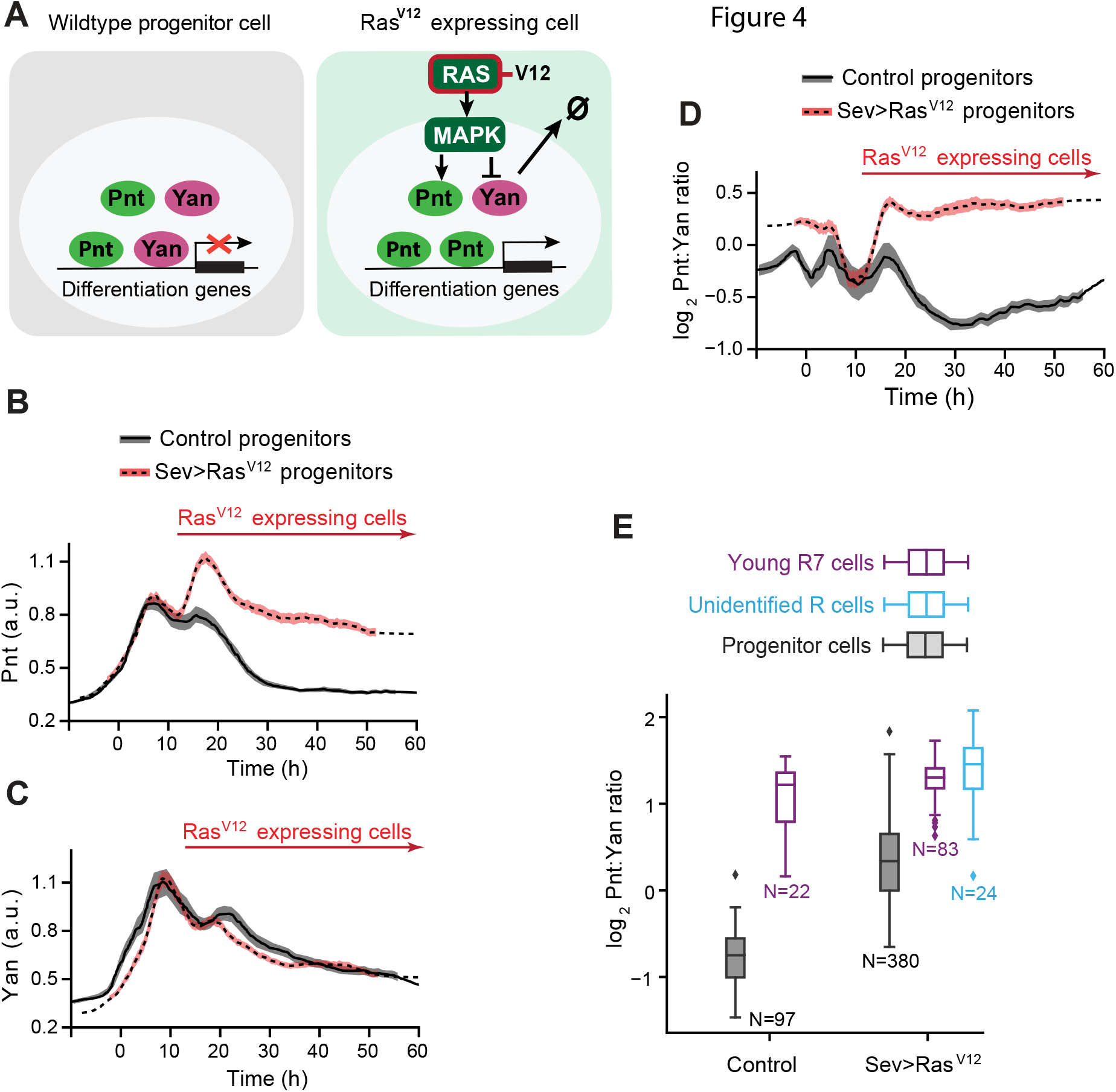
An artificial increase in the Pnt/Yan ratio biases progenitor cells towards an R fate. **(A)** The effect of the Ras^V12^ protein on signal transduction and Yan / Pnt expression in eye cells. **(B)** Line averages of Pnt-GFP protein levels in wildtype (black) and *sev>Ras*^*V12*^ (red) progenitor cells over developmental time. Shaded regions are 95% confidence intervals. Arrow marks time during which cells express Ras^V12^. For panels B – D, the number of cells analyzed in each class were 1,381 (wildtype) and 4,038 (*sev>Ras*^*V12*^). **(C)** Line averages of Yan protein levels in wildtype (black) and *sev>Ras*^*V12*^ (red) progenitor cells over developmental time. Shaded regions are 95% confidence intervals. Arrow marks time during which cells express Ras^V12^. We previously reported a modest increase in the duration of Yan-YFP expression in *sev>Ras*^*V12*^ progenitor cells (Peláez et al. 2015), but this difference was not detected using the Yan antibody. **(D)** Line averages of log_2_-transformed Pnt/Yan ratios in wildtype (black) and *sev>Ras*^*V12*^ (red) progenitor cells over developmental time. Shaded regions are 95% confidence intervals. Arrow marks time during which cells express Ras^V12^. **(E)** The log_2_-transformed ratio of Pnt/Yan in wildtype and *sev>Ras*^*V12*^ R7 cells classified as young (purple). Cells were classified as “young” by the criteria described in Figure 3A. Progenitor cells (grey) that were present in the timespan of these young R7 cells were also analyzed. In *sev>Ras*^*V12*^ eye discs, there were supernumerary R cells of an unidentifiable type (blue). Boxplots are median and quartiles. The numbers marked by N refer to the number of cells analyzed in each group.

We next asked whether inducing modest reductions in the ratio would prevent R-cell state transitions. It was previously shown that MAPK phosphorylation of Yan strongly stimulates Yan turnover (Rebay and Rubin, 1995). If the serines and threonines targeted by MAPK are mutated in Yan, then signaling-induced Yan turnover is blocked (Figure 5A). We transiently expressed this mutant Yan^ACT^ protein in progenitor cells and R3/R4/R7 cells using the *sev* promoter. The Pnt/Yan ratio in progenitor cells was not detectably altered in *sev>Yan*^*ACT*^ eye discs (Figure 5B-D and Figure S8A-C). This was not surprising since RTK – Ras – MAPK pathway is low in progenitors, and therefore Yan^ACT^ protein is not abnormally stable in these cells. Nor was the ratio altered in R8/R2/R5/R1/R6 cells of *sev>Yan*^*ACT*^ eye discs since the *sev* promoter does not induce Yan^ACT^ expression in these cells (Figure S9A-D,G,H). However, in R3/R4/R7 cells, which both receive a strong RTK signal and express Yan^ACT^, there was a reduction in the Pnt/Yan ratio. The reduction was weak in R3/R4 cells (Figures S9E,F) but was larger in R7 cells, which had a ratio value similar to progenitor cells sampled at comparable times (Figure 5B-D). Consistent with the experimental design, the modest reduction in the Pnt/Yan ratio in R cells was primarily driven by higher levels of Yan protein (Figure S8D).

**Figure 5.**
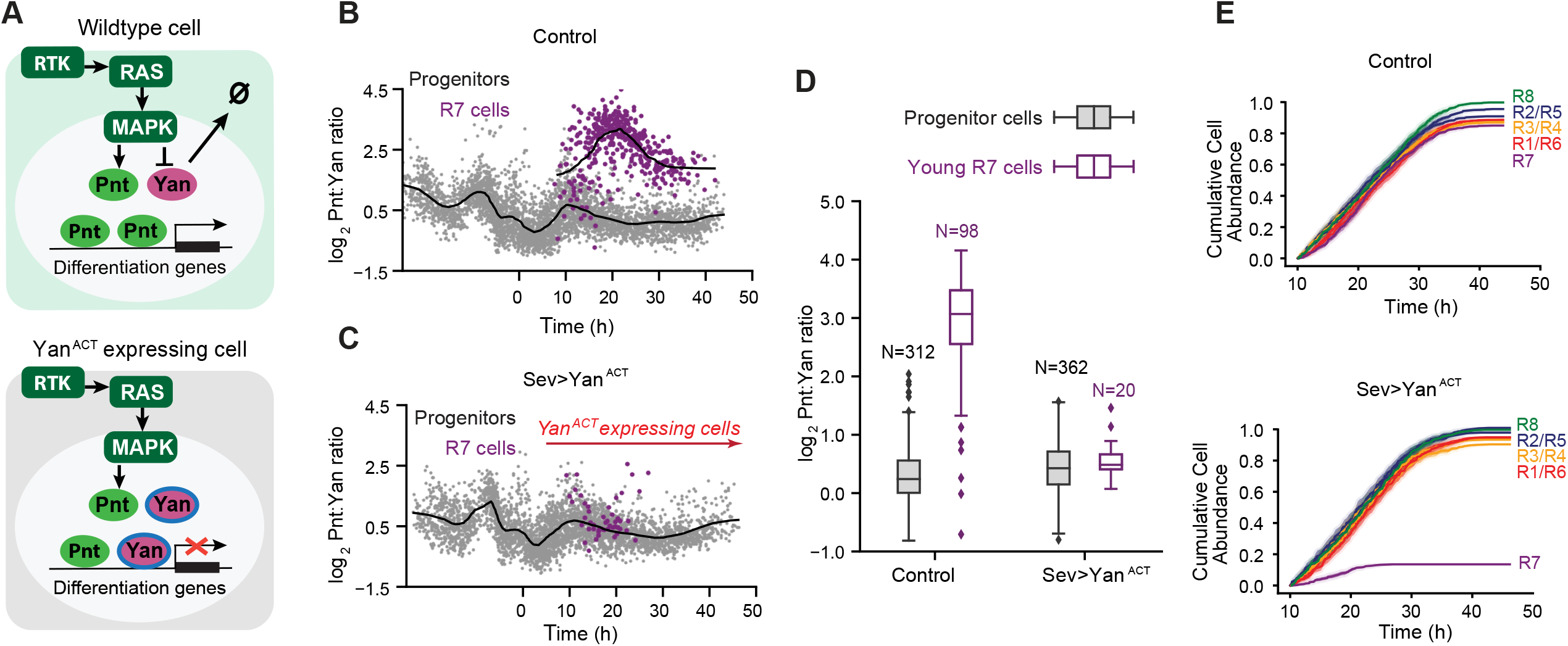
Artificial decrease in the Pnt/Yan ratio biases progenitor cells against an R fate. **(A)** The effect of the mutant Yan^ACT^ protein on signal transduction and Yan / Pnt expression in eye cells. **(B)** The log_2_-transformed Pnt/Yan ratio in wildtype control progenitor (grey) and R7 (purple) cells and their moving line averages (solid black lines). The number of cells analyzed in each class were 3,796 (progenitors) and 413 (R7 cells). **(C)** The log_2_-transformed Pnt/Yan ratio in *sev>Yan*^*ACT*^ progenitor (grey) and R7 (purple) cells and their moving line averages (solid black lines). **(D)** The log_2_-transformed ratio of Pnt/Yan in wildtype-control and *sev>Yan*^*ACT*^ R7 cells classified as young (purple). Cells were classified as “young” by the criteria described in Figure 3A. Progenitor cells (grey) that were present in the timespan of these young R7 cells were also analyzed. Boxplots are median and quartiles. The numbers marked by N refer to the number of cells analyzed in each group. **(E)** Cumulative abundance of each annotated R cell type as a function of developmental time in wildtype (top) and *sev>Yan*^*ACT*^ (bottom) eye discs. Counts are normalized by the total number of R8 cells detected across the time-course, and R2, R5, R3, R4, R1, and R6 cells are each tallied separately. Colors denote R cell type. Shading denotes 95% confidence intervals.

Changing the Pnt/Yan ratio to make it Yan-biased in young R7 cells induced a remarkable reduction in the total number of R7 cells that developed (Figure 5B,C and Figure S7A,C). Only a few R7 cells could be identified shortly after fate specification, and more mature R7 cells could not be found. Overall, the progressive fate specification of the R7 cells was reduced seven-fold in the *sev>Yan*^*ACT*^ eye discs (Figure 5E). In contrast, R3/R4 cell fate specification was not impaired even though the Pnt/Yan ratio was slightly lower than normal (Figure 5E). This suggests that cell state transitions can withstand weak or transient variation in the ratio, but not sustained larger changes.

In conclusion, two independent approaches to artificially alter the Pnt/Yan ratio resulted in cells making errors in state transitions. Modestly increasing the ratio caused cells to undergo ectopic state transitions while decreasing the ratio caused cells to fail to undergo transitions. The results suggest that progenitor cells can sense a two-fold change in the Pnt/Yan stoichiometric ratio and respond by transitioning to R-cell fates.

### A biophysical model explains how small changes in stoichiometric ratio can have a large effect on Yan and Pnt DNA occupancy

The effect of transcription factors on their target genes is highly correlated with their occupancy on DNA elements within these genes. When transcription factors do not compete for binding sites, occupancy is driven by total nuclear concentration of the transcription factor and its binding affinity. In systems with transcription factors that mutually repress each other’s expression, the stoichiometry of the factors can be very different in cells. Thus, DNA occupancy by antagonistic transcription factors reflects the large difference in their relative abundance. Since Pnt and Yan have overlapping sequence specificity for DNA binding both in vitro and in vivo, they are thought to compete for common binding sites in target genes (Flores et al., 2000; Halfon et al., 2000; Lachance et al., 2018; Nitta et al., 2015; Webber et al., 2018; Wei et al., 2010; Xu et al., 2000; Zhu et al., 2011). However, the differences in Pnt/Yan stoichiometry that we measured in eye cells are quite modest – two-fold. How could such a modest difference in stoichiometry profoundly affect relative DNA occupancy by Yan and Pnt to alter transcription of target genes?

Measuring DNA occupancy at single cell resolution is not yet experimentally feasible, so we instead investigated this question *in silico*. We first created a simple biophysical model for equilibrium binding of two transcription factors that compete for common DNA binding sites. If the proteins have a similar binding affinity for DNA and if the binding sites are saturated, then at equilibrium, occupancy will be sensitive to the relative concentration of each factor across a broad range of absolute concentrations (Figure S10).

However, the situation with Pnt and Yan is more complex. Yan binds to common binding sites with higher affinity than Pnt (Xu et al., 2000). Moreover, recent experiments suggest Yan and Pnt differentially interpret the structural syntax of enhancers and promoters (Lachance et al., 2018). This complexity is a consequence of cooperative interactions between DNA-bound Yan molecules. Yan monomers are able to self-associate via their sterile alpha motifs (SAM), enabling DNA-bound Yan monomers to stabilize the recruitment of additional Yan monomers to adjacent sites (Hope et al., 2017; Lachance et al., 2018; Qiao et al., 2004). Since Pnt does not self-associate (Mackereth et al., 2004; Meruelo and Bowie, 2009), this Yan-Yan cooperativity will bias its competitiveness for common binding sites.

Therefore, we adapted a modeling framework developed to probe the effects of *cis*-regulatory syntax on Yan binding site occupancy (Hope et al., 2017). The model considers an ensemble of microstates, each defined by a unique configuration of vacant or Yan-bound sites. Each microstate is assigned a thermodynamic potential based on the cumulative influence of strong sequence-specific binding to consensus ETS sites, weak non-specific DNA binding, and SAM-SAM polymerization. We generalized the earlier model by incorporating Pnt as a second transcription factor that competes for occupancy of the same binding sites but does not self-associate (Figure S11A,B). Using this model, we explored how Yan-Yan cooperativity affects Pnt binding site occupancy.

We first considered a scenario in which neither Yan nor Pnt exhibit cooperativity (Figure 6A). In the absence of stabilizing SAM-SAM interactions, the landscape of overall binding site occupancy was identical to that obtained with the simple equilibrium model described earlier (Figures 6A, S10, and S11). Increasing the Pnt/Yan ratio resulted in a gradual increase in Pnt occupancy for all individual binding sites, producing a titration contour closely resembling a Michaelian curve (Figures 6C and S11C).

**Figure 6.**
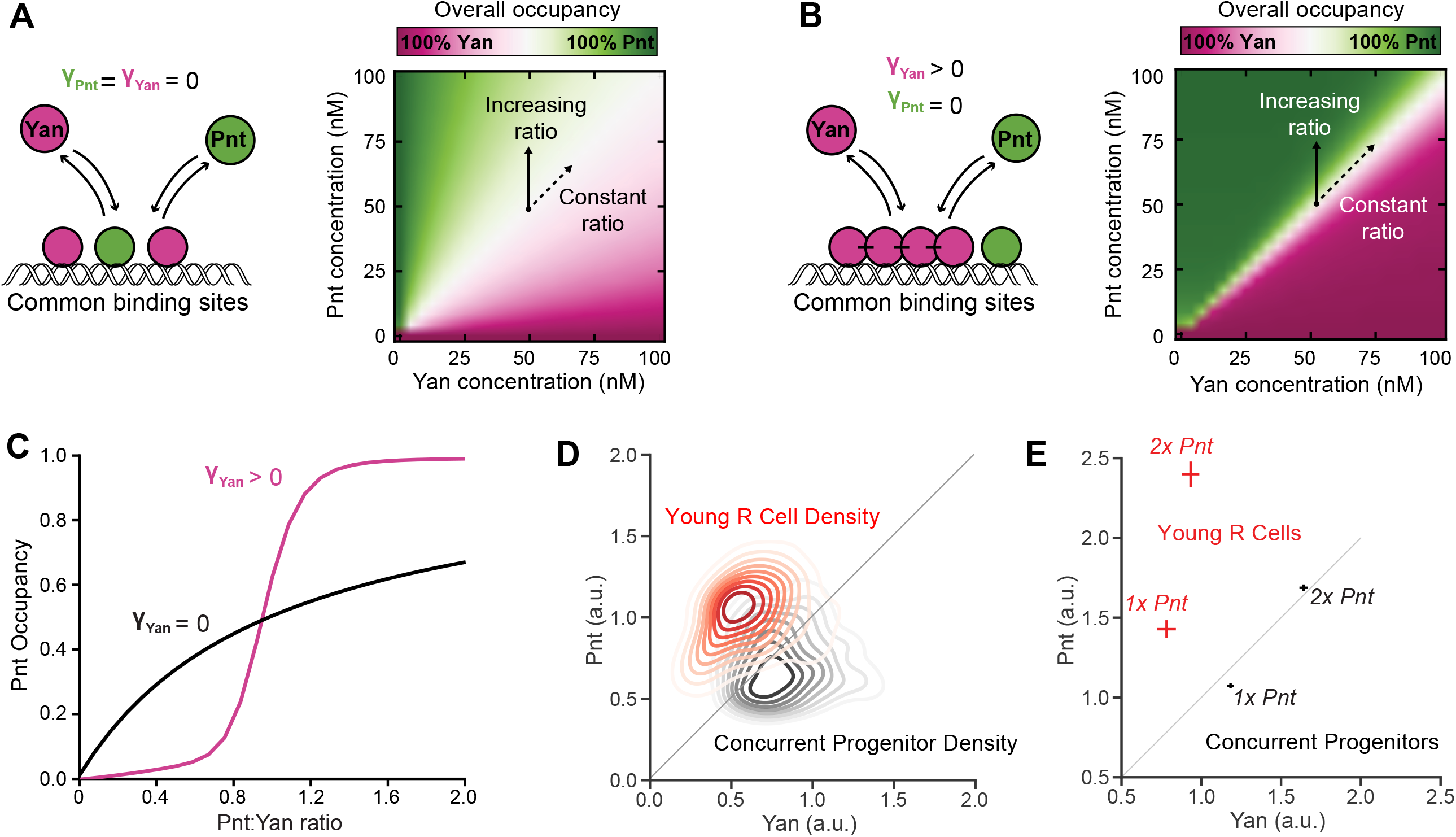
Cooperative binding by Yan greatly sensitizes DNA occupancy to the Pnt/Yan ratio. **(A)** Left – Schematic of the competition between Pnt and Yan for occupancy of mutual DNA binding sites when Yan is unable to self-polymerize. Right – Overall binding site occupancy calculated as a function of transcription factor abundance when Yan is unable to self-polymerize. We use a diverging scale because all sites are fully saturated at total transcription factor concentrations above 1 nM. Under the range of conditions shown, this implies that Yan occupies all sites left vacant by Pnt. Simultaneous proportional increases in absolute abundance of both species have minimal impact on Pnt occupancy (dashed arrow), while varying ratio confers gradual change (solid arrow). **(B)** Left – Schematic of the competition between Pnt and Yan for occupancy of mutual DNA binding sites when Yan can self-polymerize. Right – Overall binding site occupancy calculated as a function of transcription factor abundance when Yan can self-polymerize. Color scale and arrows as in (A). Note how the sharpness of the transition between all-Pnt occupancy and all-Yan occupancy is greatly increased when compared to (A). **(C)** Average Pnt occupancy across all binding sites as a function of the Pnt/Yan ratio when Yan is unable to self-polymerize (black line) versus when Yan can self-polymerize (pink line). The contours correspond to vertical paths traversed across the phase-plots in (A) and (B) at a fixed Yan concentration of 50 nM. **(D)** Probability density function of experimentally measured Pnt-GFP and Yan levels for the combined populations of all young R cells (red) and concurrent progenitor cells (black). Contours decrease in opacity with decreasing density and are not shown for values below 0.1. Density function is estimated using Gaussian kernels with a bandwidth selected using Scott’s rule. Thin diagonal line shows a constant Pnt/Yan ratio to aid interpretation. **(E)** Experimentally measured average Pnt-GFP and Yan levels for the combined populations of all young R cells (red) and concurrent progenitor cells (black), sampled from eye discs containing either one (*1x Pnt*) or two (*2x Pnt*) copies of the *pnt-GFP* gene. Horizontal and vertical lines about each point denote the standard errors of the mean. Thin diagonal line shows a constant Pnt/Yan ratio to aid interpretation.

We then introduced stabilizing SAM-SAM interactions for Yan, which dramatically sharpened the differences between Yan and Pnt occupancy (Figure 6B). The model output resembles a phase diagram in which there is a sharp boundary that separates the state in which all sites are bound by Yan from the state in which all sites are bound by Pnt. This phase boundary lies along a region in which Pnt and Yan are in 1:1 stoichiometry, regardless of total protein concentration. Weighting the energetic contributions of binding strength and polymerization by the statistical frequency of each microstate revealed that the transition was driven by an abrupt change in the dominant binding mechanism. Polymerization-associated cooperativity effects dominate binding site occupancy when the Pnt/Yan ratio is low, while binding strength dominates when the ratio is high (Figure S11D).

Increasing the Pnt/Yan ratio revealed nonlinear transitions from low to high Pnt occupancy for each individual binding site (Figures 6C and S11E). These transitions resembled Hill function forms, with sharp thresholds delimiting distinct outputs, suggesting that transitions from low to high Pnt occupancy are ultra-sensitive to changes in the ratio. At low Pnt/Yan ratios, Yan occupancy dominated, while at some critical ratio, Pnt-bound molecules prevented Yan-Yan interactions, ultimately allowing Pnt to outcompete Yan as the ratio increased further. If transcription factor levels are sufficient to saturate binding sites, then relative occupancy by Pnt and Yan was agnostic to changes in the absolute abundance of either factor, as long as the ratio remained constant.

This model suggests a molecular mechanism for how even a two-fold change in the Pnt/Yan ratio could shift DNA occupancy and therefore regulation of common target genes from an all-Yan to an all-Pnt regime. Therefore, the model provides guidance on how to interpret the experimental data. Although our fluorescence measurements do not provide us with absolute concentration estimates for Yan and Pnt, nevertheless the Pnt/Yan ratio measured in progenitor and R cells can be viewed as a parallel representation to the phase spaces calculated in the model (Figure 6B,D). In this way, the small changes in the Pnt/Yan ratio we measured could elicit large changes in DNA binding site occupancy, while significant changes in absolute Pnt and Yan concentration would have little effect so long as the relative ratio remains constant (Figure 6E). Together this ratio-based mechanism provides both ultrasensitivity and robustness to retinal cell state transitions.

## DISCUSSION

The *Drosophila* retina has an extraordinary track record as a tractable genetic system for elucidating the conserved molecular mechanisms that direct cell state transitions. By introducing single-cell resolution quantitative analysis of Yan and Pnt expression dynamics, and then computationally modeling how they might impact DNA binding complexes and hence transcriptional output, our study has uncovered important and unanticipated mechanistic features of this well-studied system. We find that eye cells in the progenitor pool maintain a dynamically stable Pnt/Yan ratio predicted to favor the Yan-dominated enhancer occupancy necessary to sustain the progenitor state. In transitioning R cells, RTK signaling disrupts the progenitor ratio set-point and produces a sustained increase in the ratio to now favor Pnt DNA occupancy. In this way, transcriptional programs associated with the progenitor state are shifted to those appropriate to the R-cell state. We suggest that sensitivity to small but sustained relative changes makes this ratiometric system robust to fluctuations in absolute Pnt and Yan protein levels while enabling rapid ultrasensitive responses to inductive signaling.

For a transition to occur, the Pnt/Yan ratio set point must be shifted. However, prior to undergoing a state transition, the progenitor’s ratio set-point is dynamically stable over time even while cells vary in Yan and Pnt expression over a four-fold range. This suggests a stabilizing mechanism that monitors the ratio of Pnt and Yan and takes corrective action when the ratio deviates from a specified reference value. The results of experiments in which we manipulated *pnt* or *yan* gene copy number suggest the stabilizing mechanism includes a weak positive feedback loop between Pnt and Yan. As a result of this positive feedback, any fluctuation in the level of one factor will bring about a compensatory change in the other factor, maintaining dynamic but approximately constant stoichiometry of the two transcription factors. Overall, this allows cell fate transitions to occur reliably over a wide range of absolute protein concentrations and prevents transient fluctuations in either Yan or Pnt from inappropriately triggering or blocking differentiation.

Our results show that experimental manipulation of Ras and MAPK activity can change the Pnt/Yan ratio, most likely by regulating the synthesis and degradation of Pnt and Yan (Graham et al., 2010). Presumably, the ratio shift that naturally occurs at state transitions is due to inductive signals coming from R8 cells, since the signals are received by RTKs and transduced by the Ras – MAPK pathway. The ratio adopts a new set-point in R cells, twice the value of the old set-point, and is sustained thereafter. If the positive feedback loop between Yan and Pnt regulates the set-point, then the strength of their mutual interactions must be differentially altered in order for the expression ratio to maintain a new value. Alternatively, feedback might remain the same but new regulatory inputs could tune Yan and Pnt expression to the new ratio in R cells. Investigating the contributions of the multiple regulatory interactions that have been documented within the Pnt/Yan network (Graham et al., 2010; Rohrbaugh et al., 2002; Webber et al., 2013b; Webber et al., 2018; Wu et al., 2020) to maintenance and modulation of the ratio will be an interesting future direction.

Previous studies have shown how cells can sense relative concentration changes of individual molecular species using fold-change detection (Frick et al, 2017, Adler & Alon, 2018, Alon et al., 1999; Barkai and Leibler,1997; Cohen-Saidon et al., 2009; Goentoro and Kirschner, 2009; Lee et al., 2014; Mesibov et al.,1973; Shoval et al., 2010). To compute fold-change of a single protein, regulatory circuits first “measure” and establish a molecular memory of the state of one variable, and later compare it to its background level or to a new measurement of the same variable (Frick et al, 2017, Lyashenko et al., 2020). In contrast, rather than measuring the individual fold-change in Pnt or Yan over time, eye cells likely rely on concurrent comparison of Pnt and Yan levels, or more speculatively as suggested by the biophysical model, by concurrent comparison of the transcriptional outcome of Yan and Pnt competition for DNA binding sites.

Progenitor cells exhibit pulsatile expression of Yan and Pnt in which their expression levels decay to a basal state by 50 hours of developmental time. If cells transit to a R-cell fate, this decay in expression is accelerated. Nevertheless, the Pnt/Yan ratio remains stable about a fixed set-point specific for cells in either progenitor or R-cell states. This is in contrast to many developmental systems in which the ratio of antagonistic transcription factors remains stable because expression of the factors is maintained constant when cells exist in a particular developmental state (Laslo et al., 2008). A ratiometric system with modest variation may provide several advantages not just for the eye, but potentially to other systems.

First, such a system can permit fast response times. In the eye, a cascade of four inductive events occurs over a brief 8-hour window of time to recruit R1 – R7 cells. If a classical mechanism of mutual transcriptional repression existed between Yan and Pnt expression, then the time to resolve the ratio change would likely be too slow. By using a system in which a two-fold shift in the ratio triggers cells to switch states, transient RTK inputs are sufficient to change the relative levels of the two proteins, thereby robustly and rapidly achieving the objective.

Second, a ratio-sensing system can define precise temporal windows of competence for differentiation. In the eye, competence of cells to respond to RTK signals is not only dependent on the transduction machinery but also on the presence of Yan and Pnt, which regulate target gene transcription downstream of RTKs. Eye progenitor cells anterior to the MF do not express Yan and Pnt, likely to prevent these cells from inappropriately responding to RTK activity. Over a 50-hour time window as the MF moves across the eye disc, progenitor cells undergo the first phase of cell fate specification during which R cells and cone cells are recruited into ommatidia. Since fate specification occurs as a wave across the eye, posterior ommatidia complete the first specification program ∼50 hours before anterior ommatidia do. Yet posterior progenitor cells do not carry on with further fate transitions until the anterior ommatidia complete the initial induction of R cells and cone cells (Cagan and Ready, 1989). It is only then, that progenitor cells resume undergoing RTK-induced fate transitions – forming the pigment cells (Freeman, 1996; Miller and Cagan, 1998). We suggest that Yan and Pnt are shut off after the R-cell specification phase in order to create a multi-hour gap in progenitor competence that separates the two phases of fate specification.

Finally, a ratio-based mechanism can confer ultrasensitivity to modest changes in molecular stoichiometry when it is coupled to additional molecular mechanisms, for example, competition and cooperativity in enhancer occupancy. Although the ratio of Pnt and Yan shifts only by two-fold when cell state transitions occur, this change is sufficient to produce robust regulation of the transit to differentiation. Our biophysical model shows how the composition of transcription factor complexes on Yan/Pnt target genes could dramatically change when a two-fold ratio change occurs, going from a repressive to an activating composition. Stabilizing interactions between Yan molecules bound to adjacent DNA sites sensitize enhancer occupancy to the relative abundance of the competing transcription factors. Indeed, prior work has shown that DNA-bound Yan monomers enhance recruitment of Yan to adjacent binding sites through cooperative protein-protein interactions (Lachance et al., 2018; Qiao et al., 2004). Given that cooperativity and competition are features of many biological systems, ratiometric control mechanisms analogous to the one we have described in the Drosophila eye could prove to be a broadly used regulatory strategy.

## EXPERIMENTAL PROCEDURES

### Genetics

Unless otherwise noted, all experiments were performed with animals reared at 25°C on standard cornmeal-molasses *Drosophila* culture medium. Fly stocks from the Bloomington Stock Center: *pnt-GFP* BAC transgene, BL42680; *pnt*^*2*^, BL2222; H2Av-mRFP BL23650, pnt^Δ88^ (O’Neill et al., 1994); *sev>Ras*^*v12*^ (Fortini et al., 1992); *sev>Yan*^*ACT*^ (Rebay and Rubin, 1995); *sev>Gal4* (Jemc and Rebay, 2006). Measurements of wild type dynamics of Pnt-GFP were made in eye discs from *w*^*1118*^; *pnt-GFP; pnt*^*Δ88*^*/pnt*^*2*^, *H2Av-mRFP*. The effect of *pnt* gene dosage on Pnt/Yan ratio was measured in *w*^*1118*^; *pnt-gfp /+* ; *pnt*^*Δ88*^*/pnt*^*2*^ (1x *pnt*) and *w1118; pnt-gfp; pnt*^*Δ88*^*/pnt*^*2*^ (2x *pnt*) discs. In experiments with *Ras*^*v12*^ or *Yan*^*ACT*^, Pnt-GFP and Yan levels were measured in eye discs from animals of the following genotypes: *w*^*1118*^; *pnt-gfp/ sev>Ras*^*v12*^; *pnt*^*Δ88*^*/+* relative to *w*^*1118*^; *pnt-gfp / +; pnt*^*Δ88*^*/+*; and *w*^*1118*^; *pnt-gfp/sev>Yan*^*ACT*^; *pnt*^*Δ88*^*/+* relative to *w*^*1118*^; *pnt-gfp/ +; pnt*^*Δ88*^*/+*. Yan mutant eye clones were generated using the *yan*^*833*^ null allele, *ey>FLP* and the FRT40 crossover point. *Yan*^*+*^ cells were labeled using the clonal marker *Ubi>mRFP*_*nls*_ (Bloomington Stock 34500). Developing eyes were dissected from white prepupae carrying *w, ey>FLP* ; *pnt-gfp, yan*^*833*^, *FRT40A/ pnt-gfp, Ubi>mRFP*_*nls*_, *FRT40A*.

### Immunohistochemistry

All eye-antennal discs were dissected from white prepupae. For experiments in which 4′,6-diamidino-2-phenylindole (DAPI) was used as the nuclear marker, samples were fixed in 4% paraformaldehyde (w/v) / PBS-Triton X-100 0.1% (v/v) for 25 minutes at room temperature. After fixation, eye discs were blocked in PBS-Triton X-100 0.1% (v/v) and 1% (v/v) normal goat serum for 30 minutes at room temperature. Primary and secondary antibodies were incubated each for 2 hours at room temperature with DAPI. Including DAPI during primary and secondary antibody staining was important for even penetration of DAPI. Antibodies used: mouse anti-Yan 8B12 (DHSB, 1:200) and goat anti-mouse Cy3 (1:2000, Jackson Immunoresearch). Discs were washed for 20 min in PBS-Triton X-100 0.1% (v/v), and mounted in VectaShield Antifade mounting medium (Vector Laboratories). To avoid rapid changes in cell volume, which would affect relative fluorescence measurements, the substitution of PBS-Triton X-100 with Vectashield was done gradually using increments of ∼33% in VectaShield concentration. For experiments in which H2Av-mRFP was used as the nuclear marker, discs were fixed in 4% (w/v) paraformaldehyde/PBS for ∼45 min and goat anti-mouse Pacific Blue antibody (Life Technologies, 1:200 dilution) was used as the secondary antibody.

Samples were imaged within 18-24 hr after fixation. 1024 × 1024 16-bit images were captured using either a Zeiss LSM880 or a Leica SP5 confocal microscope equipped with 40X oil objectives. Discs were oriented with the equator (dorsal-ventral boundary) parallel to the x-axis of the image. Optical slices were set at 0.8μm (45-60 optical slices per disc) with an additional digital zoom of 1.2-1.4. Images recorded a region of at least 6 rows of ommatidia on each side of the equator. All discs for a given perturbation were fixed, mounted, and imaged in parallel with control discs to reduce measurement error. Sample preparation, imaging, and analysis were not performed under blind conditions, nor was sex of the animals noted at time of dissection.

### Quantification of expression levels

A minimum of three experimental replicate eye discs were imaged for each experimental condition, with the exception of the *sev>Ras*^*V12*^ condition for which only one control disc was used. For each wild type eye disc, ∼160 R8, ∼320 R2/5, ∼300 R3/4, ∼260 R1/6, ∼ 220 R7 and >1000 progenitor cells were manually selected from a set of computationally 2D-segmented nuclei using Silhouette. The fluorescence intensity of each nucleus was measured by our established procedure (Peláez et al., 2015). Our new pipeline includes Silhouette, an open-source package for macOS that integrates our image segmentation algorithm with a GUI (graphical user interface) for cell type annotation (available at: https://www.silhouette.amaral.northwestern.edu/). Subsequent analysis and visualization procedures were carried out using the accompanying FlyEye python package, which is freely available on GitHub. The resulting expression dynamics were inferred from confocal image stacks using an updated version of the segmentation and annotation pipeline described in (Peláez et al., 2015). Empirically, the replicate number of three eye discs provided reproducible composite dynamics. No outlier eye discs were found for any experimental condition, so all samples were pooled, temporally aligned and included in the analysis. The sample size was not pre-computed before data collection.

Nuclear segmentation was performed using either H2Av-mRFP (Figures 1, 3, 4, S1-S2, S4 and S5) or DAPI (Figures 2, 5 and S7-S9) as a reference channel to identify individual nuclear boundaries. Nuclei expressing H2Av-mRFP were segmented using the Silhouette app’s optimized implementation of the GraphCut algorithm (QI, 2013), while DAPI-stained discs were segmented using an open-source implementation of the watershed algorithm designed to mitigate the effect of bright spots caused by DAPI accumulation in nucleoli (Bernasek et al., 2020). Each layer of the confocal stack was segmented in 2D independently in both cases. For each nucleus measured, a single contour was manually selected and assigned a cell type (cell state) using Silhouette, with care taken to avoid annotating nucleolar contours rather than complete nuclear contours in DAPI-stained discs. Since each nucleus spans multiple optical sections, contours were always selected in the middle of the nucleus. The population of progenitor cells was measured in multiple layers to include cells whose nuclei were at different positions in the Apical - Ventral axis of the eye disc. In all cases each nucleus was measured only in one layer to avoid measurement pseudoreplication.

Cells were identified and annotated by nuclear position in xyz space and shape using the His2Av-mRFP channel with the other channels turned off. marker protein channel turned off. To validate the accuracy of this method of cell identification / annotation, cells were also immunostained for various cell-type specific marker proteins, as noted in Table S1. Annotations made with tie His2Av-mRFP channel were then compared to the cell-type specific markers visualized in other channels.

For each annotated nucleus contour, expression measurements were obtained by normalizing the mean fluorescence of the Pnt-GFP and Yan antibody channels by the mean fluorescence of the reference channel used for segmentation (DAPI or His2Av-mRFP). This normalization mitigates variability due to potentially uneven sample illumination, segment area, nuclear volume, and in the case of His-RFP, differences in protein expression capacity between cells.

### Conversion of distance to time

Cell positions along the anterior-posterior axis were mapped to developmental time as described previously (Peláez et al., 2015). This is predicated on two assumptions: the furrow proceeds at a constant rate of one column of R8 neurons per two hours; and minimal cell migration occurs. For each disc, Delaunay triangulations were used to estimate the median distance between adjacent columns of R8 neurons (Fortune and Steven, 1992). We used the median rather than the mean distance because it minimized the influence of non-adjacent R8s that were falsely identified by the triangulation (Peláez et al., 2015). Dividing the furrow velocity of 2 h per column by this median distance yields a single conversion factor from position along the anterior-posterior axis to developmental time. This factor was applied to all cell measurements within the corresponding disc, yielding expression time series. Notably, these are not single cell dynamics, but rather aggregate dynamics across the development time course of a spatially organized cell population.

### Computation of moving averages and confidence intervals

Moving averages were computed by first-order Savitzky-Golay filtration (Savitzky and Golay, 1964). This method augments the simple windowing approach used in (Peláez et al., 2015) by enabling visualization of expression trends at early time-points that are otherwise obscured by large window sizes. A secondary first-order filtration with one-fifth the original window size was applied to smooth lines for visualization purposes. None of our conclusions are sensitive to our choice of filtration or smoothing method. Primary window sizes of 250 and 75 cells were used for reporting the expression of multipotent and differentiated cells, unless noted otherwise. Confidence intervals for the moving average were inferred from the 2.5th and 97.5th percentile of ten-thousand-point estimates of the mean within each window. Point estimates were generated by bootstrap resampling with replacement of the expression levels within each window.

### Alignment of expression data

To align multiple eye disc samples, cells of each sample were aligned with a reference population by shifting them in time. The magnitude of this shift was determined by maximizing the cross-correlation of progenitor Pnt-GFP expression *Y(t)* with the corresponding reference time series *X(t)*. Rather than raw measurements, moving averages within a window of ten cells were used to improve robustness against noise. This operation amounts to:

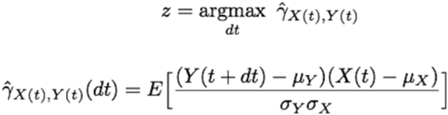

where, *μ* and *σ* are the mean and standard deviation of each time series, and *dt* is the time shift by which the population should be shifted.

For each experimental treatment, a disc was randomly chosen and shifted in time such that time zero corresponds to the first annotated R8 neuron. This disc then served as the reference population for the alignment of all subsequent biological replicates within the treatment. Similarly, different experimental treatments (e.g. control and perturbation) were aligned by first aligning the discs within each treatment, then aggregating all cells within each treatment and repeating the procedure with the first treatment serving as the reference.

This approach differs from the previous implementation of our pipeline in which discs were manually aligned by the inflection point of their Yan-YFP expression profiles (Peláez et al., 2015). Manual alignment entails arbitrary prioritization of certain dynamic features over others. Our revised protocol yields consistent, reproducible alignment of expression time series that equally weighs the entire time course. The automated approach is more principled but less robust than the manual approach.

Specifically, it fails when dynamic forms qualitatively differ between experimental treatments. In these instances, we revert to manual alignment using the inflection point of Pnt-GFP induction as a reference.

### Analysis of *yan* clones

We used *ey>FLP* and *FRT40A* to generate *yan*^*833*^ null clones within 23 eye discs carrying the Pnt-GFP transgene (see Genetics section). The chromosome carrying the wildtype *yan* allele was marked with a Ubi>mRFP_nls_ transgene. Discs were dissected, fixed, and co-stained with DAPI prior to confocal imaging of RFP and GFP. Images of 36 unique vertical cross-sections spanning non-overlapping cells were collected in total. Images were analyzed using our previously published method called Fly-QMA that quantitatively analyzes mosaic imaginal discs (Bernasek et al., 2020). Briefly, cell nuclei were identified by watershed segmentation of the DAPI channel. For each segment, Ubi-mRFPnls and Pnt-GFP fluorescence was quantified by normalizing the average intensity of all pixels within the respective fluorescence channel by the average DAPI fluorescence. Fluorescence bleed-through between the GFP and RFP channels was corrected as described previously (Bernasek et al., 2020). Bleed-through correction successfully eliminated any detectable difference in Pnt-GFP expression in mosaic eye discs in which all cells were wildtype for *yan*. The same correction procedure was therefore applied to all measurements of mosaic eye discs containing *yan* mutant clones.

Mitotic recombination between the mutant and wildtype chromosomes yields cell populations exhibiting low, medium, and high levels of RFP fluorescence, which correspond to cells with 0,1, and 2 copies of the wildtype allele, respectively (Bernasek et al., 2020). Segmented nuclei were assigned to one of three groups using a k-means classifier. The procedure was validated through comparison with manual annotation of ∼2,500 cells. The overall classification rate was ∼95%, Cells residing on the border of clones were excluded from all analyses to mitigate edge effects. The remaining measurements were aggregated across all eye discs for comparison of cells with zero versus one copy of the wildtype *yan* gene.

### Simple competitive binding model

Figure S10 presents results for an equilibrium model of two species, Yan (Y) and Pnt (P), competing for a finite pool of shared binding sites, S:

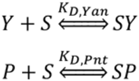

where K_D,Yan_ and K_D,Pnt_ are equilibrium association constants and SY and SP denote the bound species. Applying a mass balance to the total protein and binding site (S_0_) abundances:

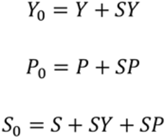

yields an analytically tractable system of nonlinear equations (Wang, 1995). For each pair of absolute protein abundances (Y_0_, P_0_), the Pnt binding site occupancy is simply SP/S_0_.

### Competitive binding model with cooperativity

The model presented in Figures 6 and S11 expands upon the work of (Hope et al., 2017). The model is based on a single cis-regulatory element consisting of *n* adjacent binding sites, each of which

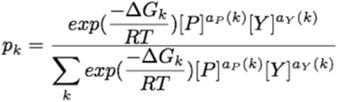

may be designated as ETS or non-ETS. Each binding site may exist in one of three binding states; bound by a single copy of Yan, bound by a single copy of Pnt, or unbound. Thermodynamic potentials were assigned to each binding state using two parameters for each transcription factor. The parameter *αX* defines the free energy of transcription factor *X* binding to an ETS site, while *βX* defines the free energy of binding to a non-ETS site. A unique configuration of binding states for all *n* binding sites constitutes a single microstate, *k*. The thermodynamic potential of each microstate was taken to be the sum of thermodynamic potentials for each of its constituent binding sites. For each microstate, the stabilizing effect of polymerization was incorporated via a third parameter, *γX*, that defines the free energy of SAM-SAM binding between a pair of similar transcription factors bound to adjacent sites. The net result is a total thermodynamic potential, *ΔGk*, for each microstate. An example enumeration of all possible microstates for an element consisting of one ETS site preceding two non-ETS sites is provided in Figure S11B. The statistical frequencies of each microstate were evaluated by constructing a canonical ensemble: in which *pk* is the statistical frequency of microstate *k, [P]* is the Pnt concentration, *[Y]* is the Yan concentration, *aP(k)* and *aY(k)* are functions representing the number of bound molecules of *P* and *Y* within microstate k, *T* is a fixed temperature set to 300 K, and *R* is the gas constant. Fractional occupancies for each binding site correspond to the cumulative statistical frequency of all microstates in which the site is occupied by a given transcription factor. Overall fractional occupancies are similarly evaluated across all sites within the element.

We considered regulatory elements comprised of 12 binding sites in which only the first site carries the ETS designation. We retained the same parameterization of Yan binding used by (Hope et al., 2017): *αY*=-9.955 kcal mol^-1^, *βY*=-5.837 kcal mol^-1^, and *γY*=-7.043 kcal mol^-1^. We parameterized Pnt binding thermodynamics to provide balanced competition between Pnt and Yan in the absence of any SAM-mediated polymerization of Pnt. That is, we set Pnt binding affinities such that the transition from Pnt to Yan occupancy occurs when Pnt and Yan concentrations are approximately equal. The model used to generate Figure 6B assumes that Pnt binds individual sites with elevated affinities *αP* = 0.96(*αY* + *γY*) and *βP* = 0.96(*βY* + *γY*). The model used to generate Figure 6A uses these same elevated binding affinities for Yan, while setting *γY*=0 kcal mol^-1^. Qualitatively, our results are not sensitive to this parameterization.

Statistical frequencies of all microstates were enumerated using a recursive graph-traversal algorithm implemented in python, where each node in the graph represents an individual binding site and each edge reflects occupancy by a specific transcription factor. An open-source python implementation of this modeling framework has been published on GitHub (https://doi.org/10.5281/zenodo.5520493).

### Data and software availability

All segmented and annotated eye discs will be deposited in a data repository upon publication.

Data are deposited at https://datadryad.org/stash/share/FrDw4tWe_ddlP7ATp3oYEvLG6FMe_3XAvNkbp9Ma2fk. A series of Jupyter notebooks that use these data to reproduce all analysis, simulations, and figures presented in this manuscript has also been made available (https://github.com/sbernasek/pnt_yan_ratio).

The Silhouette app used to segment nuclei in eye discs expressing His-2Av-mRFP and annotate nuclei in all eye discs is freely available via the macOS App Store, while the FlyEye python package used to generate, align, and visualize expression dynamics has been published on GitHub (https://doi.org/10.5281/zenodo.5520468).

## ACKNOWLEDGEMENTS

We thank members of the Amaral, Carthew and Rebay labs for helpful discussions during the course of this project, and C. LaBonne and M. Glotzer for helpful comments on the manuscript. We thank Kevin White’s lab and the MODEncode Consortium for recombineering the Pnt-GFP transgene, the Howard Hughes Medical Institute (HHMI) and the Hanna H Gray Fellowship program (N.P.), the Chicago Biomedical Consortium (N.P), the Robert Lurie Northwestern Cancer Center (N.P.), the NIH (EY025957, I.R. and R.W.C; EY032233, R.W.C.; GM118144, R.W.C.; GM080372, I.R.), the NSF (1764421, L.A.N.A., N.B. and R.W.C), and Simons Foundation (597491 L.A.N.A., N.B. and R.W.C) for funding, the Bloomington Drosophila Stock Center for flies, Laura Nilson for the use of computational resources, the Developmental Studies Hybridoma Bank for antibody reagents, and the Northwestern Biological Imaging Facility (BIF) for technical imaging support.

## SUPPLEMENTARY FIGURE LEGENDS

**Figure S1.**
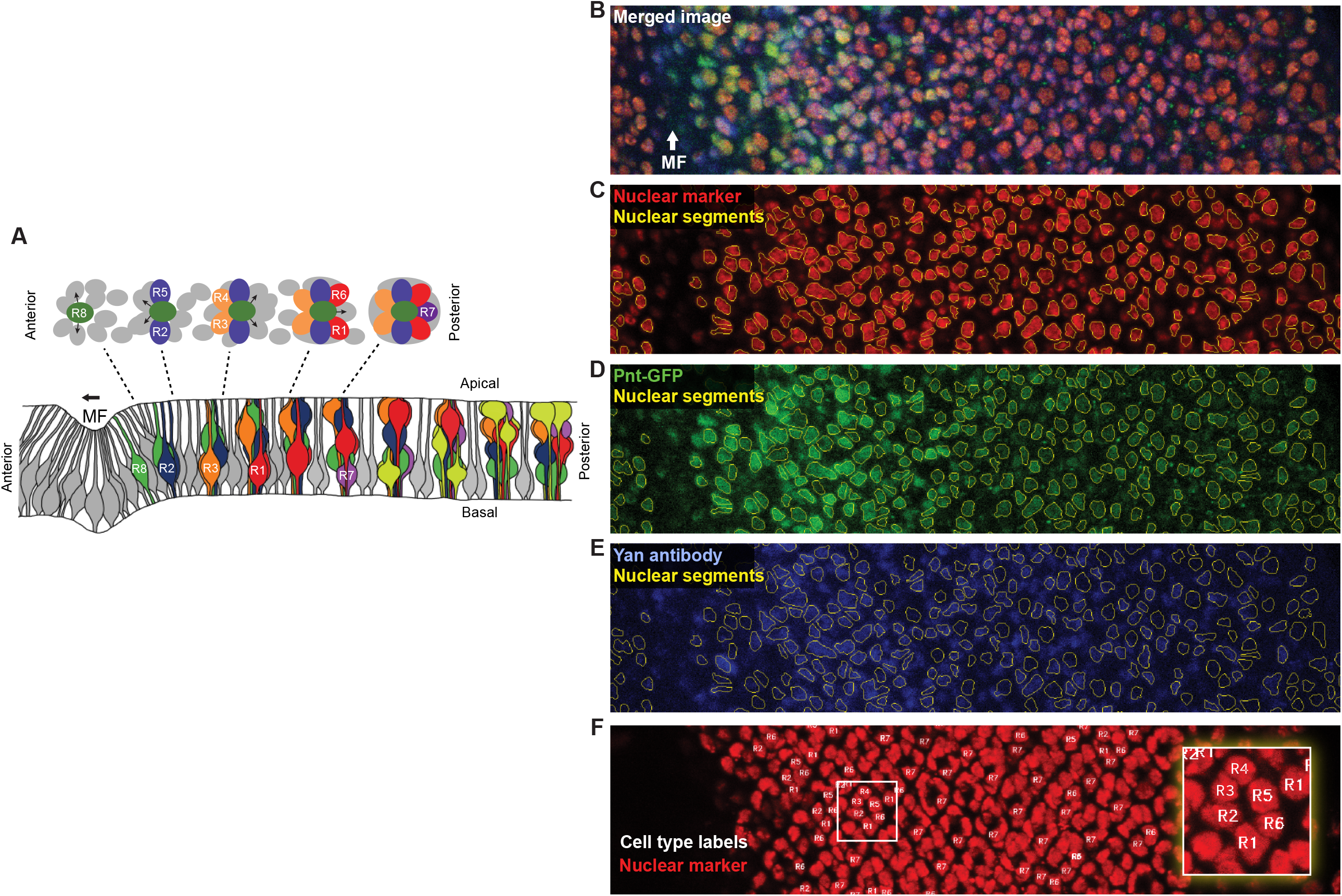
Identification of cell nuclei in eye discs. **(A)** Schematic cross-sectional view along the anteroposterior axis of an eye disc, showing the epithelial invagination that marks the MF and the assembling clusters of differentiated cells forming ommatidia. Note the dynamic and characteristic apical-basal positions of nuclei from progenitor cells (grey), R cells (various colors), and cone cells (light green). Shown above is a apical-basal view of five ommatidia sampled at different locations along the anteroposterior axis. The relative positions of progenitor and R cell nuclei are highly stereotyped in this plane. Arrows denote signals transmitted from the R8 to neighboring progenitor cells and transduced via the RTK – Ras – MAPK pathway. Adapted from (Peláez et al., 2015). **(B-E)** Quantitative analysis pipeline showing microscopy and segmentation of one representative eye disc. Shown is a partial cross-section of a confocal z-stack with the merged channels (B) and separate fluorescence channels for Histone-RFP (C), Pnt-GFP (D), and Yan antibody (E). MF marks the location of the MF. Yellow lines depict nuclear contours segmented by FlyEye Silhouette. **(F)** Manual annotation of R-cell types. Image shows a projection of His-RFP fluorescence across the Z-sections spanning R cell nuclei. The region imaged is identical to that shown in (B-E). The R-cell types are documented and are labelled as shown. The inset shows an ommatidium magnified to see nuclear positions.

**Figure S2.**
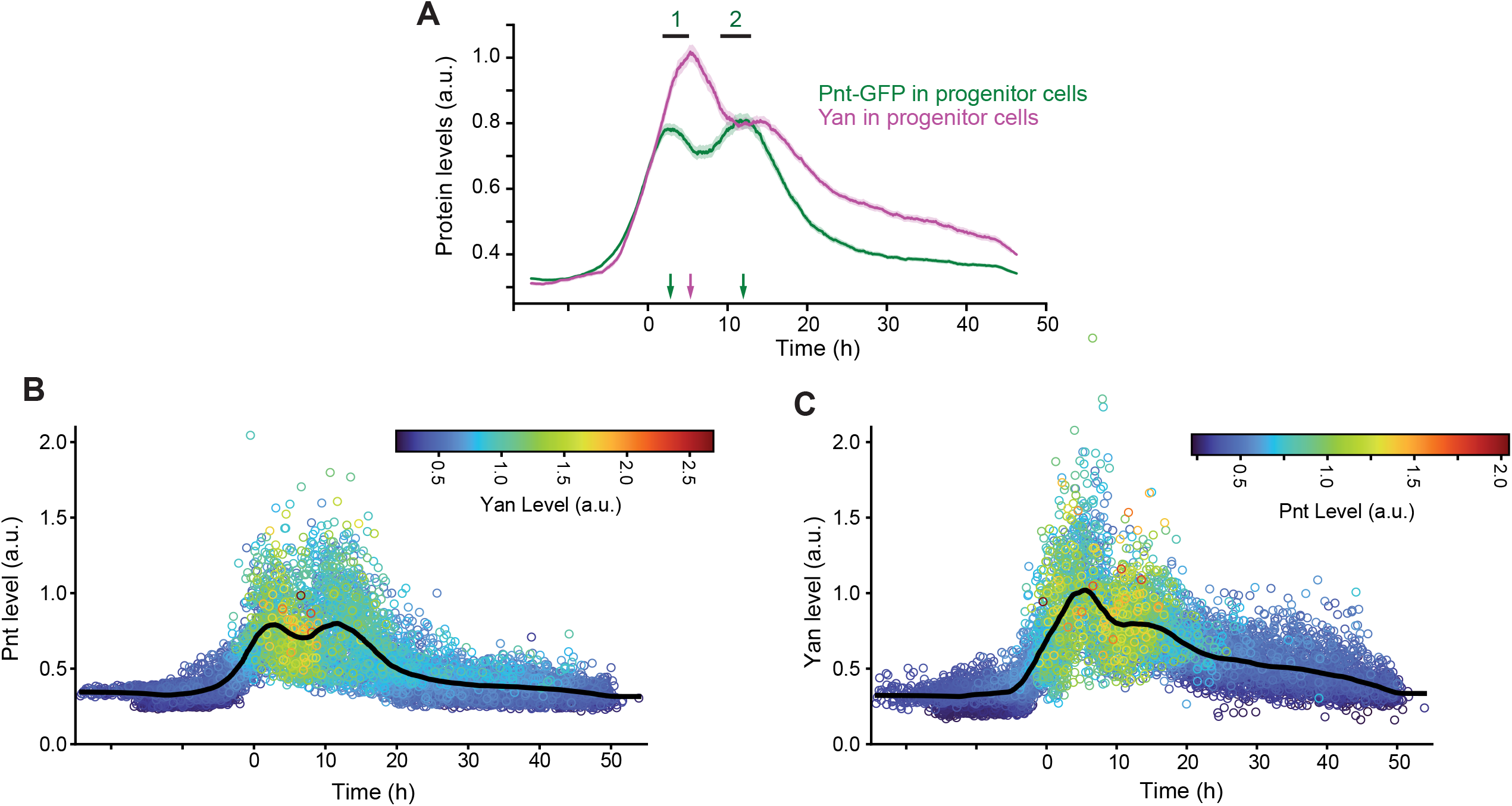
Pnt-GFP and Yan expression in progenitor cells. **(A)** Line averages of Pnt-GFP (green) and Yan (magenta) levels in progenitor cells. Lines are smoothed moving averages across 500 sequential progenitors, shaded regions are 95% confidence intervals. Arrows indicate the times at which local maxima occur. Note the peaks of Pnt and Yan expression take place sequentially in progenitor cells. **(B, C)** Expression levels of Pnt-GFP (B) and Yan (C) in individual progenitor cells, color-coded in accordance with the expression level of the complementary transcription factor, as shown at the right of each panel. Datapoints are arranged such that those with higher levels of the complementary transcription factor are displayed in front. Lines are smoothed moving averages across 500 sequential progenitor cells.

**Figure S3.**
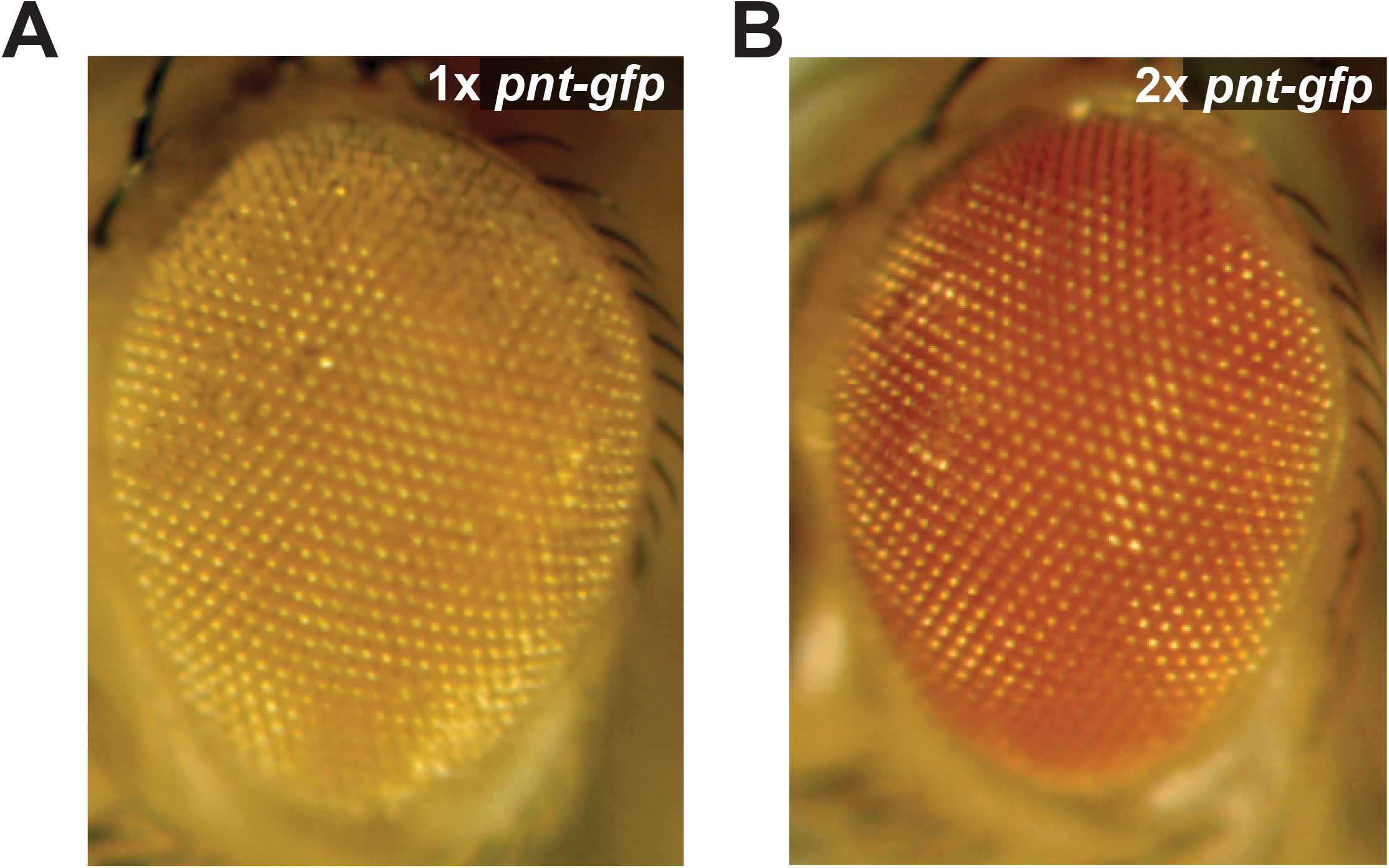
Characterization of Pnt-GFP. Adult eyes of flies carrying **(A)** one or **(B)** two copies of the *pnt-gfp* transgene in a *pnt* null mutant background.

**Figure S4.**
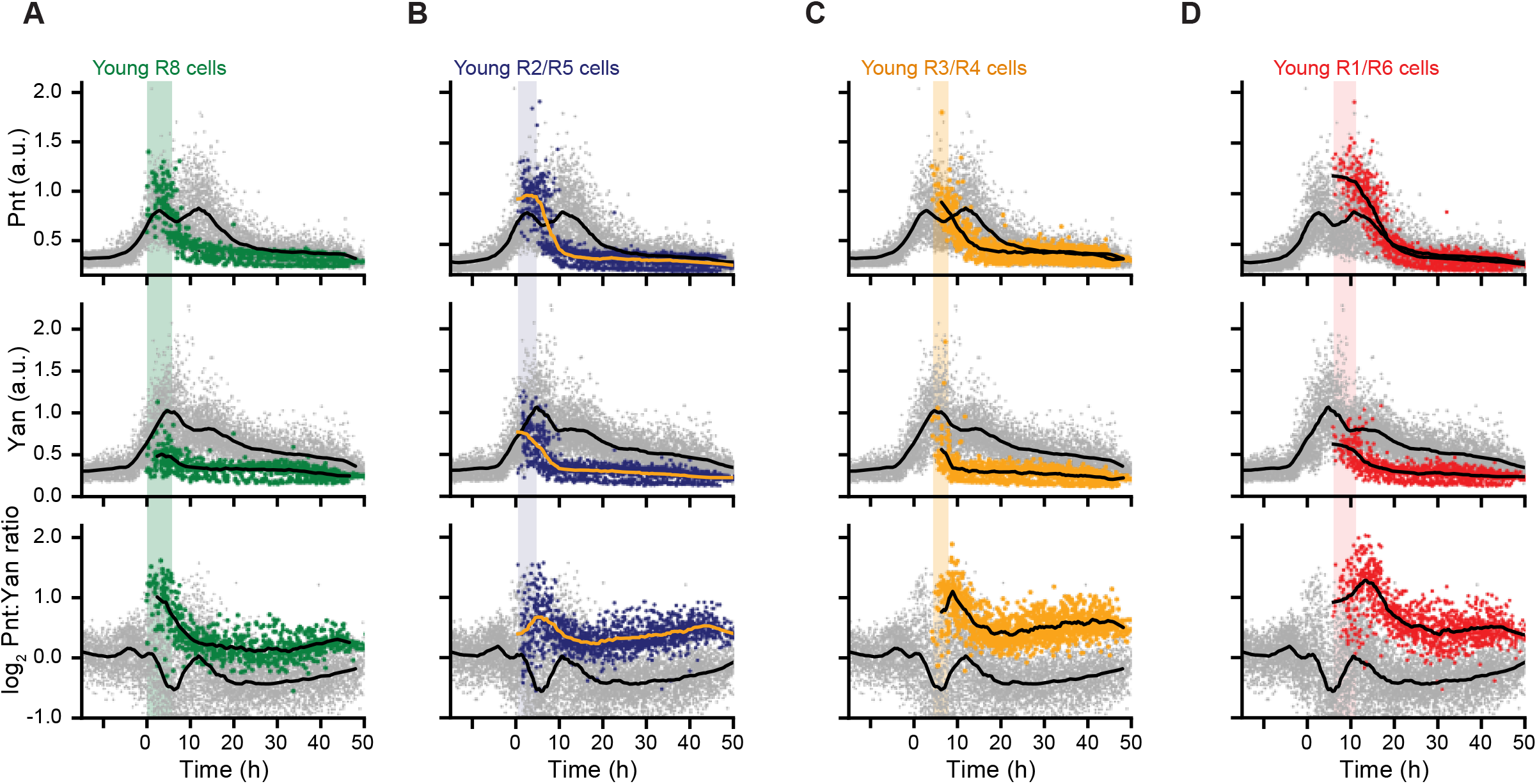
Expression dynamics for various eye cell types. **(A-D)** Pnt-GFP and Yan protein levels, and the log_2_-transformed Pnt/Yan ratio for individual progenitor (grey), and R8 cells (**A**), R2/R5 cells (**B**), R3/R4 cells (**C**), and R1/R6 cells (**D**) over developmental time. Progenitor cells are present across all time (grey arrow) while R cells arise later in time (colored arrows). Solid black lines are smoothed moving averages across 250 and 75 individual nuclei for progenitor and R cells, respectively. Black bars labeled 1 and 2 denote the two peaks of Pnt-GFP in progenitor cells. Shaded vertical stripes highlight these two regions, which coincide with transition of progenitor to various R fates.

**Figure S5.**
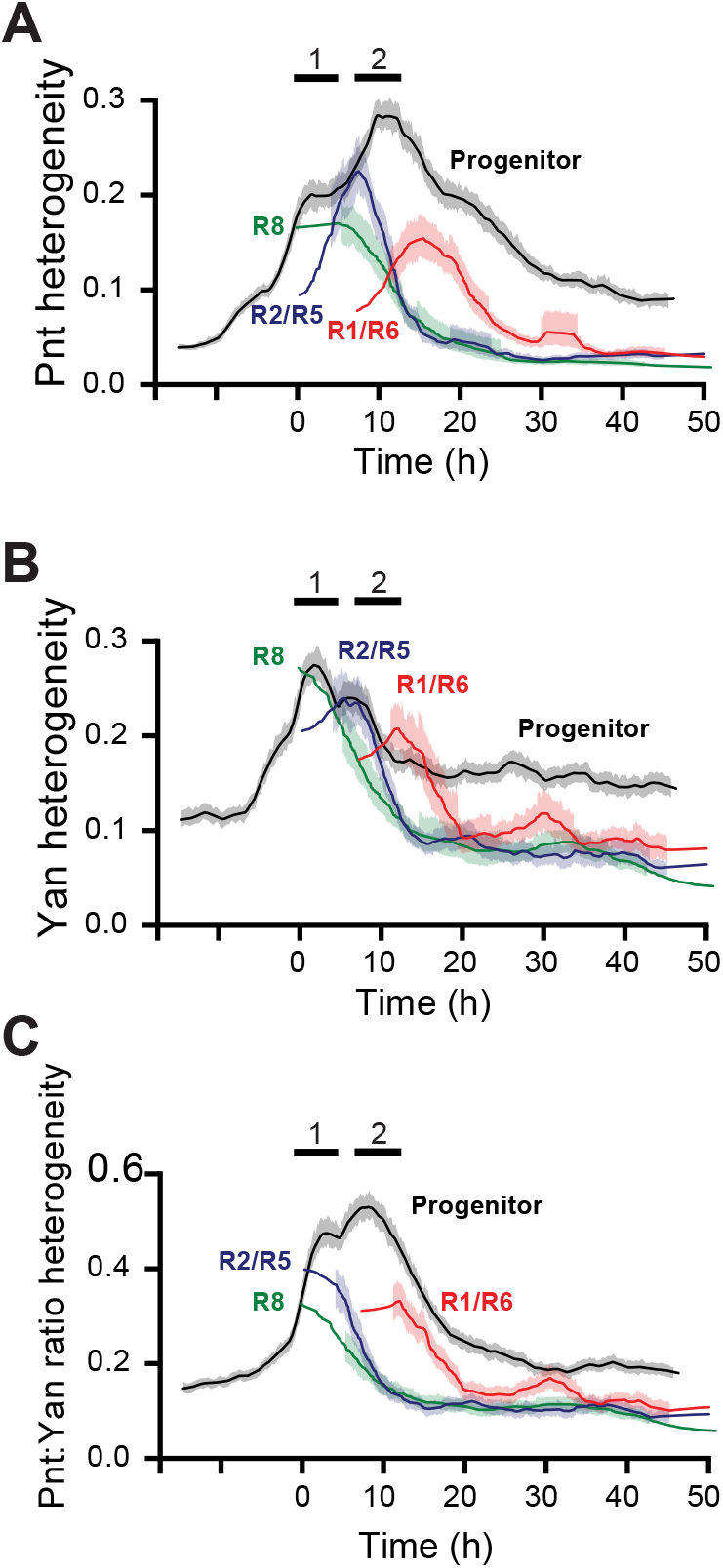
Heterogeneity of Yan and Pnt expression. Cell-to-cell expression heterogeneity of Pnt-GFP **(A)**, Yan **(B)**, and the log_2_-transformed Pnt/Yan ratio **(C)**. Heterogeneity was estimated by de-trending fluctuations about a moving average of 250 sequential cells as described (Paleaz 2015). Lines shown are moving averages of 250 sequential fluctuations. Shaded regions are 95% confidence intervals. Colors denote cell type.

**Figure S6.**
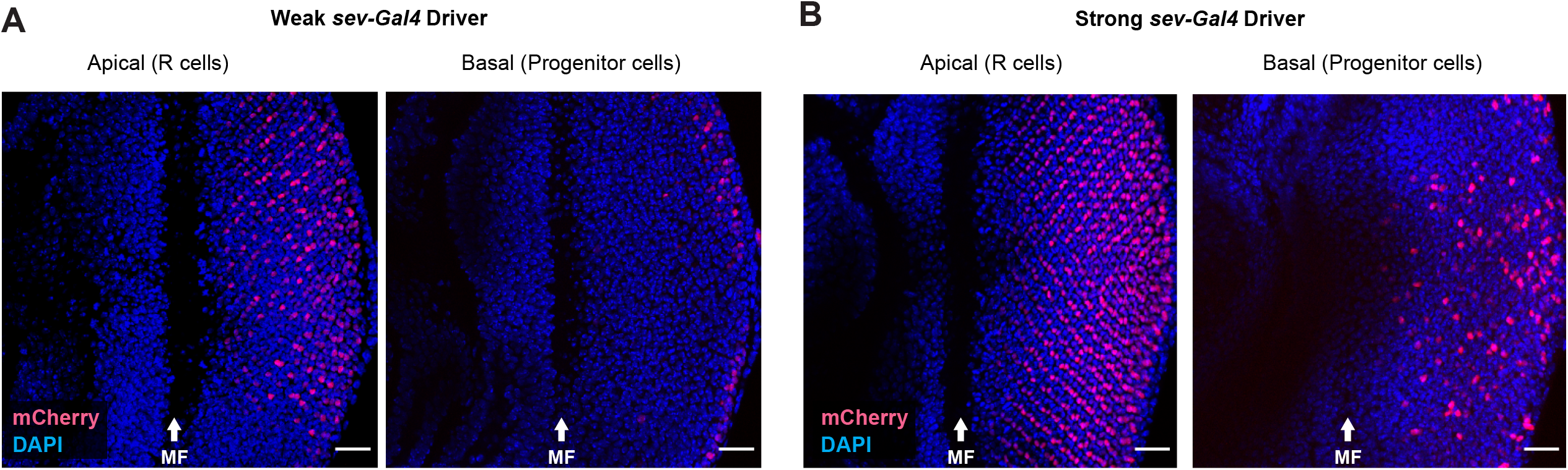
*Sev-Gal4* drives UAS transgene expression in subsets of progenitor cells. Projection of *sev-Gal4>UAS-NLS-mCherry* protein fluorescence (red) and DAPI (blue) in apical z-slices containing R-cell nuclei and basal z-slices containing progenitor cell nuclei. Anterior is left and MF is morphogenetic furrow. Scale bars denote 20 μm. **(A)** Expression from a weak *sev-Gal4* driver. **(B)** Expression from a strong *sev-Gal4* driver. When expressed from the weak driver, mCherry is predominantly limited to R7, R3, and R4 cells. When expressed from the strong driver, mCherry is also detected in some progenitor cells.

**Figure S7.**
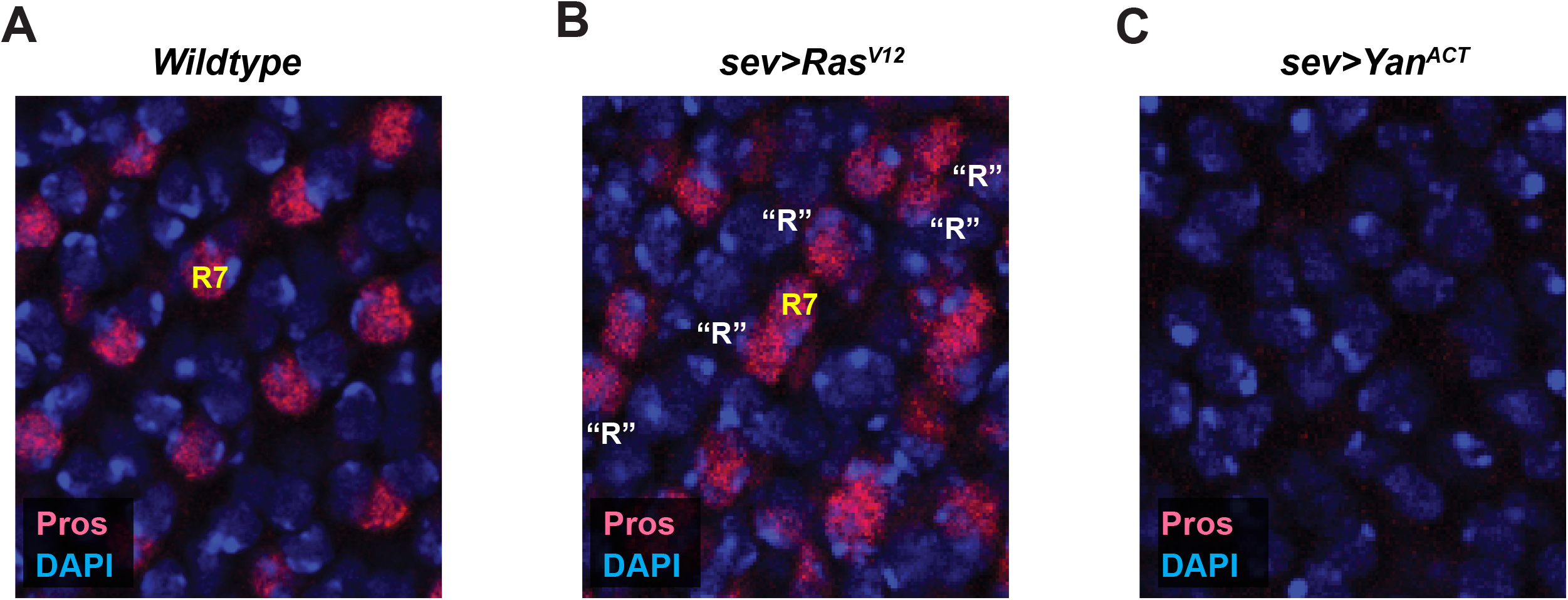
Fate specification errors in *sev>Ras*^*V12*^ and *sev>Yan*^*ACT*^. Prospero (Pros) protein fluorescence in a region of the eye disc where ommatidia have recruited cells to form R7 cells. Pros is specifically expressed in R7 cells. **(A)** Ommatidia in wildtype eye disc, which have one Pros-positive cell per ommatidium. **(B)** Some ommatidia in a *sev>Ras*^*V12*^ eye disc have supernumerary R cells labelled “R” that are Pros-positive and located beside R7 cells. **(C)** Ommstidia from a *sev>Yan*^*ACT*^ eye disc have no Pros-positive cells, indicating that they lack R7 cells.

**Figure S8.**
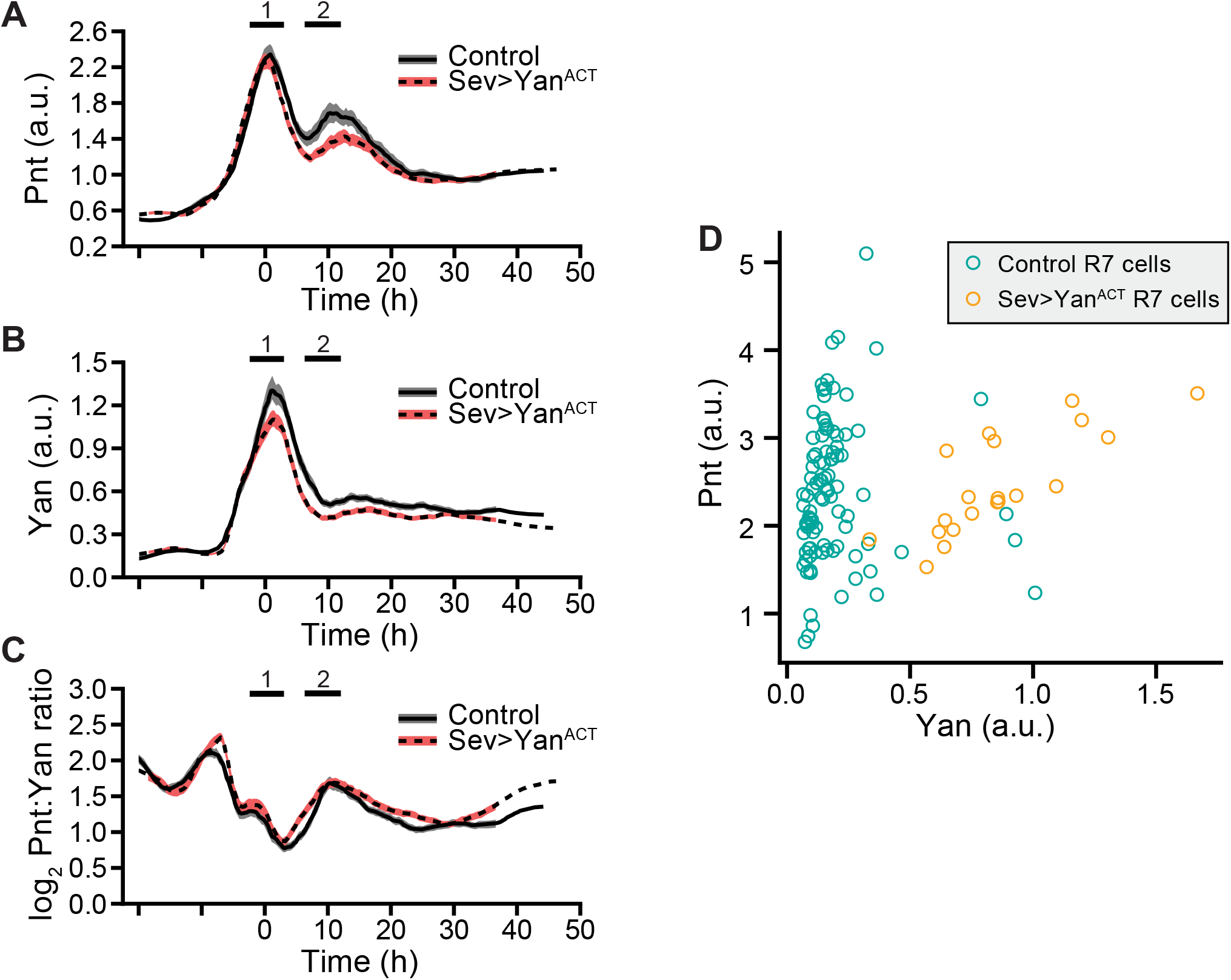
Yan^ACT^ has no effect on the Pnt/Yan ratio in progenitor cells but does alter Yan levels in R7 cells. **(A-C)** Effect of *sev>Yan*^*ACT*^ expression on Pnt-GFP (A), Yan (B), and the log_2_-transformed Pnt/Yan ratio (C) in progenitor cells over time. Lines are moving averages across 250 sequential cells. Shaded regions are 95% confidence intervals. Black lines denote wildtype cells. Red lines denote *sev>Yan*^*ACT*^ cells. **(D)** Joint distribution of Pnt and Yan levels among R7 cells sampled from the region spanning 15 to 20 h of developmental time. The *sev>Yan*^*ACT*^ R7 cells show higher Yan levels (P<0.001, KS 2-sample test), while Pnt levels are indistinguishable from wildtype (P=0.4, KS 2-sample test).

**Figure S9.**
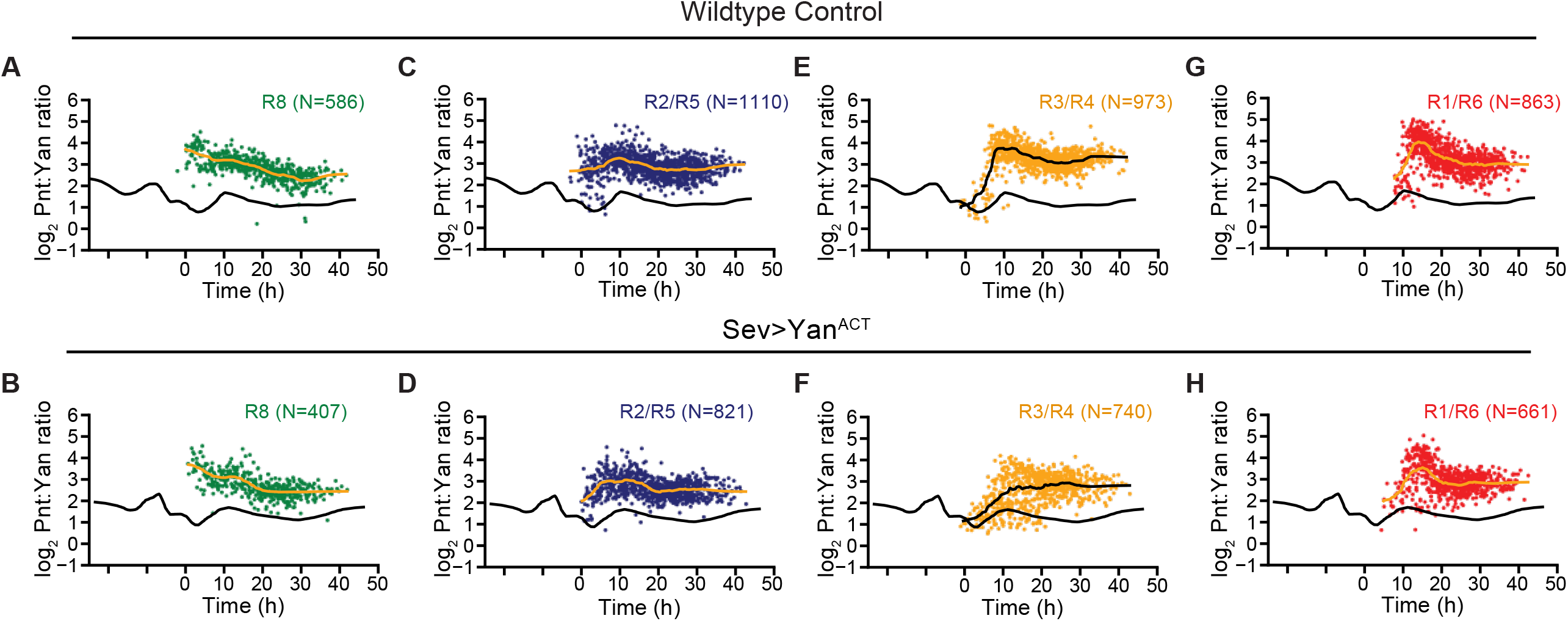
Yan^ACT^ has a weak effect on the Pnt/Yan ratio in R3/R4 cells. The log_2_-transformed Pnt/Yan ratio for individual R cells (dots) and their moving line averages (black lines) over developmental time in wildtype **(A**,**C**,**E**,**G)** and *sev>Yan*^*ACT*^ eyes **(B**,**D**,**F**,**H)**. The moving line average for progenitor cells (black lines) is also shown in each plot. Black bars labeled 1 and 2 denote the two peaks of Pnt-GFP in progenitor cells, which coincide with transition of progenitor to various R fates. Various R cell types are shown, R8 **(A**,**B)**, R2/R5 **(C**,**D)**, R3/R4 **(E**,**F)**, and R1/R6 **(G**,**H)**.

**Figure S10.**
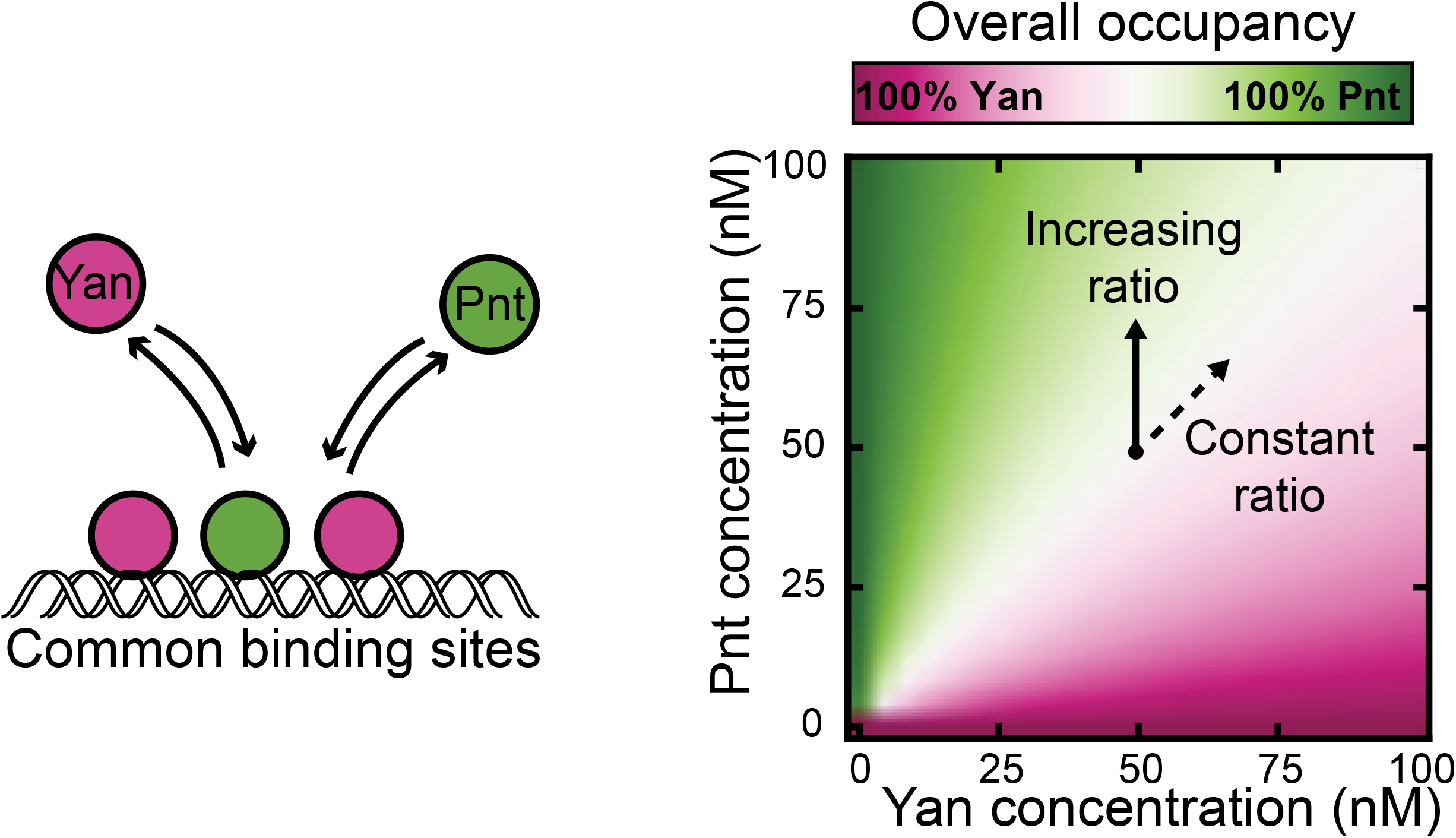
A simple equilibrium model of Yan / Pnt competition for DNA binding sites. Schematic of a simple two-species competitive binding model (left), alongside theoretical Pnt site occupancy as a function of transcription factor abundance (right). Equivalent binding affinities are used for illustrative purposes. Color scale reflects overall Pnt site occupancy. A diverging scale was used because all sites are fully saturated at total transcription factor concentrations above 1 nM. Under the range of conditions shown, this implies that Yan occupies all sites left vacant by Pnt. Simultaneous proportional increases in absolute abundance of both species have minimal impact on Pnt occupancy (dashed arrow), while varying ratio confers gradual change in occupancy (solid arrow).

**Figure S11.**
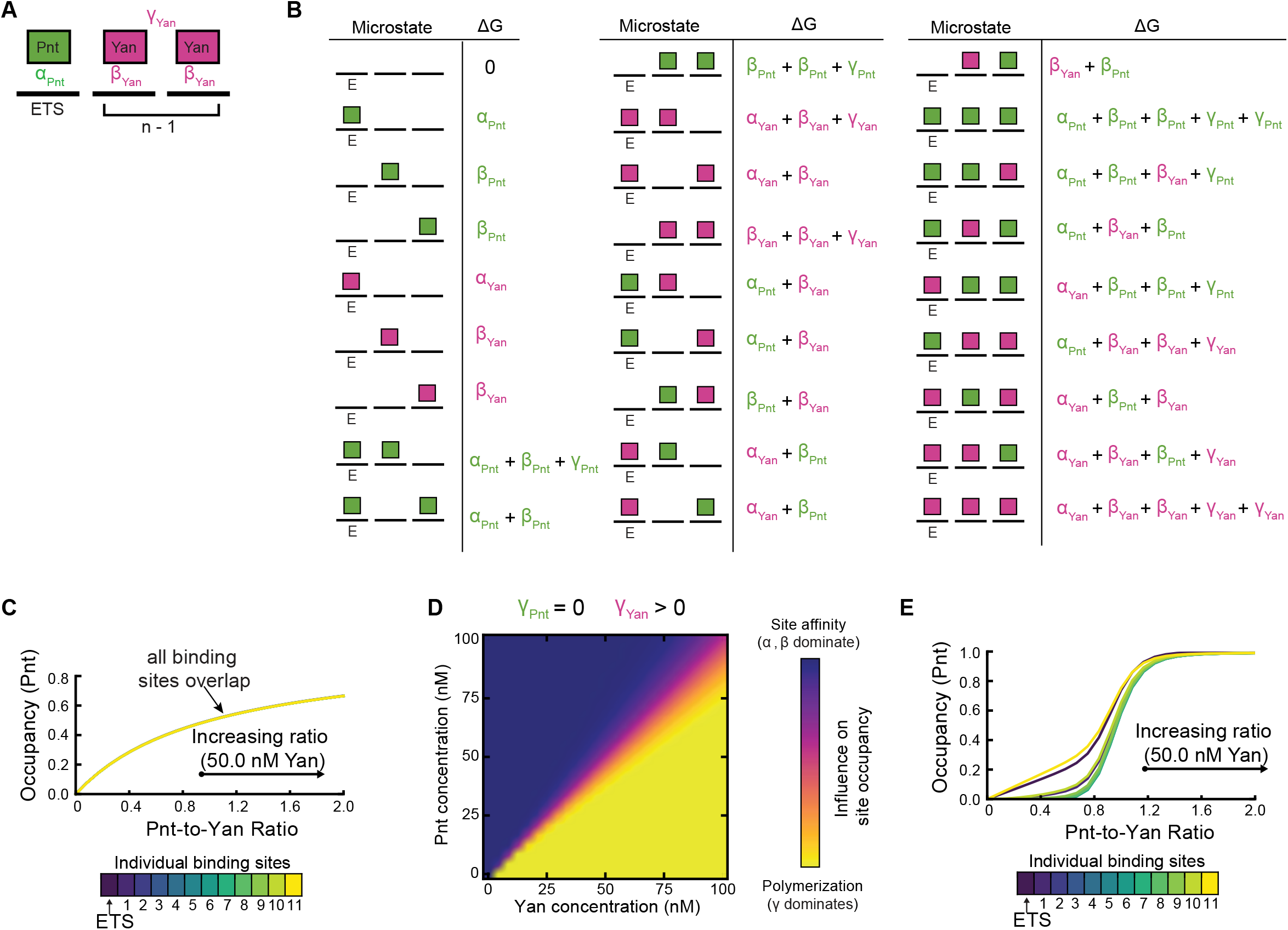
Cooperative DNA-binding by Yan sensitizes occupancy of DNA sites to the Pnt/Yan ratio. **(A)** Summary of thermodynamic interactions within one microstate of a cis-regulatory element containing one ETS site and two non-ETS sites. Solid black lines represent individual binding sites. Green and magenta rectangles denote Pnt and Yan molecules. Example thermodynamic potentials of strong ETS-binding, weak non-ETS binding, and polymerization interactions are denoted by *αPnt, βYan*, and *γYan*, respectively. For this microstate, *aP(k)*=1 and *aY(k)*=2. **(B)** Enumeration of all possible microstates for a cis-regulatory element of length 3 in which only the first site carries the ETS designation. Solid black lines denote binding sites, green and magenta rectangles denote bound Pnt and Yan molecules. The cumulative thermodynamic potentials of each microstate, *ΔGk*, are listed beside each graphical depiction. **(C)** Pnt occupancy of individual binding sites as a function of Pnt/Yan ratio in the absence of Yan polymerization. Contours correspond to a vertical path traversed across Figure 6A at a fixed Yan concentration of 50 nM. All binding sites behave similarly. **(D)** Relative thermodynamic contributions of binding site affinity versus polymerization to microstate statistical frequencies as a function of Pnt and Yan concentration. For each point in the plane, the influence of site affinity was calculated by weighting the sum of all ETS and non-ETS thermodynamic potentials for each microstate by the statistical frequency of the corresponding microstate. The influence of polymerization was similarly determined. The color scale reflects the relative magnitude of these two summations, normalized by limits of zero and complete polymerization. **(E)** Pnt occupancy of individual binding sites as a function of Pnt/Yan ratio when Yan polymerizes. Contours correspond to a vertical path traversed across Figure 6B at a fixed Yan concentration of 50 nM. Line colors reflect binding site positions within the cis-regulatory element. Sites at intermediate distances from the strong ETS site (green lines) transition at higher ratios than those nearest and furthest from the strong ETS site (blue and yellow lines).

**Table S1.** Accuracy of cell-type identification

| **Cell Type Scored** | **Cell Type Marker** | **Sample Size (*n*)** | **% Correct Type Assignment** |
| --- | --- | --- | --- |
| **Progenitors** | Elav (-) | 462 | 99.8 |
| **All mature Rs** | Elav (+) | 258 | 99.2 |
| **R8** | Sens | 106 | 97.2 |
| **R2/R5** | Rough | 162 | 99.4 |
| **R3/R4** | Rough | 142 | 100.0 |
| **R7** | Pros | 104 | 99.0 |

